# Structural Dynamics and Allosteric Communication of a SARS-Like Bat Coronavirus Spike Glycoprotein

**DOI:** 10.64898/2025.12.12.693972

**Authors:** Toheeb A. Balogun, Fiona L. Kearns, Carla Calvó-Tusell, Alexandra L. Tse, Cory M. Acreman, Lorenzo Casalino, Gorka Lasso, Emily Happy Miller, Kartik Chandran, Jason S. McLellan, Rommie E. Amaro

**Affiliations:** Department of Molecular Biology, University of California, San Diego, La Jolla, California, United States of America; Institute of Physical Chemistry, University of Freiburg, Albertstrasse 21, 79104 Freiburg, Germany; Department of Microbiology & Immunology, Albert Einstein College of Medicine, Bronx, New York, New York, United States of America; Department of Molecular Biosciences, The University of Texas at Austin, Austin, Texas, United States of America; Division of Biology and Biological Engineering, California Institute of Technology, Pasadena, California, United States of America; Division of Infection and Immunity, University College London, London, UK; Department of Medicine, Albert Einstein College of Medicine, Bronx, New York, New York, United States of America

**Keywords:** SHC014, bat coronavirus, allosteric communication, molecular dynamics simulations, disease spillover

## Abstract

SARS-like bat coronaviruses (CoVs) pose ongoing public health risks due to their zoonotic potential, making it important to understand the molecular pathways driving their evolution. We recently showed that SHC014-CoV can infect human cell lines in an ACE2-dependent manner after acquiring two spike ectodomain mutations (F294L and A835D). However, how the wild-type (WT) SHC014 spike differs dynamically from these mutants remains unclear. Here, we built fully glycosylated ectodomain models of WT and three mutants (F294L, A835D, and the double mutant, DM) and performed triplicate 1-μs all-atom molecular dynamics (MD) simulations for each variant. The two mutations exhibit epistasis, altering structural rearrangements relative to WT. Notably, the DM receptor binding domain (RBD) begins sampling the open conformation in our conventional MD. At the atomic level, the DM spike mitigates the dense negative packing introduced by A835D through a salt-bridge network, while F294L disrupts π-mediated interactions, together enhancing RBD opening propensity—critical for viral entry. Increased flexibility of the subdomain-2 “620-loop” further modulates DM RBD openness. Dynamical network analysis identified three allosteric communication pathways. In WT and F294L, “Pathway 1” forms the baseline route linking the 620-loop to the RBD, whereas in A835D and DM it extends to the FPPR, reshaping long-range communication. “Pathway 2” is conserved across variants but is most prominent in WT and F294L. “Pathway 3” appears only in A835D and DM, compensating for reduced communication along Pathway 2. Overall, this work provides an atomistic perspective on SHC014 molecular adaptation during host-to-host transmission and highlights mechanistic features that may inform future therapeutic and pandemic-preparedness efforts.

**Statement of Significance:** Bat coronaviruses are an important source of future pandemic threats, but we still know little about how small genetic changes help them infect humans. In this study, we used detailed computer simulations to watch how tiny mutations in a bat coronavirus spike protein change its motion and shape. We found that two specific mutations work together to make the spike more likely to open—a step required for the virus to enter human cells. By revealing how these molecular changes increase infection potential, our work helps improve understanding of coronavirus evolution and may guide strategies to prepare for future outbreaks.

## Introduction

Bats harbor SARS-like coronaviruses (SL-CoVs) with zoonotic potential that continue to pose emerging viral threats to human health.^1^ There have been multiple outbreaks of coronaviruses, including severe acute respiratory syndrome coronavirus (SARS-CoV), Middle East respiratory syndrome coronavirus (MERS-CoV), and SARS-CoV-2, with their origins hypothesized to stem from bats.^2,3^ Several SL-CoVs have been identified^4–7^ in nature such as RatG13, RpYN06, BANAL-52, RmYN02, WIV1, that share close genome identity with SARS-CoV-2, the human infectious agent of the COVID19 pandemic. Furthermore, various studies have shown that the spike protein in certain bat coronaviruses can utilize the human angiotensin-converting enzyme 2 (ACE-2) and its orthologs for cell entry, thereby priming SL-CoV to mediate infections in humans.^8–12^

A metagenomics study identified seven strains of SL-CoVs from Chinese horseshoe bats in Yunnan Province, China, including two novel pre-emergent strains called RsSHC014 and Rs3367, and elucidated their full-length genome sequences.^7^ Although the authors were unable to recover a replication-competent RsSHC014, hereafter referred to as SHC014, they were able to characterize the first replication-competent SL-CoV, WIV-1-CoV, which is closely related to Rs3367. A follow-up study by Menachery and colleagues^13^ showed that pseudotyped SHC014 is unable to utilize the human cellular receptor ACE2. Surprisingly, chimeric virus containing the SHC014 spike glycoprotein was found to be replication competent by using multiple orthologs of the ACE2 receptor, can infect human airways, and cause disease in mice, depicting SHC014’s zoonotic potential to cause spillover events from bats to other mammals, including humans.

In SARS-CoV, SARS-CoV-2, and SHC014, the spike glycoprotein (GP) (**Figure 1**) is a class 1 fusion homotrimer protein that covers the surface of coronaviruses and plays a key role in viral entry.^14,15^ Furin cleaves the transmembrane spike protein at the polybasic recognition site (PRRAR) known as the furin cleavage site (FCS). Cleavage at the FCS splits the spike sequence into two subunits, the S1 and the S2.^16–21^ The S1 subunit consists of the N-terminal domain (NTD) and a receptor binding domain (RBD), which contains the receptor binding motif (RBM), the region that binds directly to ACE2 and mediates host cell entry.^22,23^ S2 is responsible for fusion of viral and host cell membranes and is comprised of the fusion peptide (FP), two heptad repeats (HR1 and HR2), a transmembrane domain (TM), and a palmitoylated cytoplasmic tail (CT) (**Figure 1A**).^15,24^ Several studies have elucidated the structures of the SARS-CoV-2 spike protein in various conformational states.^25–28^ Such SARS-CoV-2 spike conformational states are characterized by the number and degree of “up” or exposed RBDs: the closed spike conformation being one in which all three of the spike RBDs are “down” or highly shielded, and 1-up, 2-up, and 3-up states indicate conformations with one, two, or three up RBDs respectively^27,29,30^. In the down state, the RBM within an RBD is highly shielded by neighboring protein surfaces and N-linked glycans, thus down RBDs are considered to be inaccessible for ACE2 binding.^31^ In an up or open states (more pronounced up states), the RBM is unshielded and can bind to ACE2.^27,30–32^ During RBD opening, several other domains are known to also move including the NTD, two S1 subdomains (SD1 and SD2), the Fusion Peptide Proximal Region (FPPR), as well as several glycans(**Figure 1A,1B**).^17,33,34^ Glycans are carbohydrate-based post-translational modifications anchored either to asparagines (N-linked) or serines/threonines (O-linked) that cover the spike GP, aiding in host-cell immune evasion^25,26,35–38^. Our group has previously elucidated the structural role of several N-linked glycans, particularly those at positions N165 and N234, in stabilizing the human SARS-CoV-2 spike’s RBD in the up position.^31^ Similarly, we have previously described the gating role of the N343 glycan in facilitating RBD opening.^30^ Therefore, interactions between spike subunits and glycans can modulate the conformational plasticity of spike RBDs, impacting accessibility to the ACE2 host cell receptor and therefore driving membrane fusion and viral entry.

**Figure 1:**
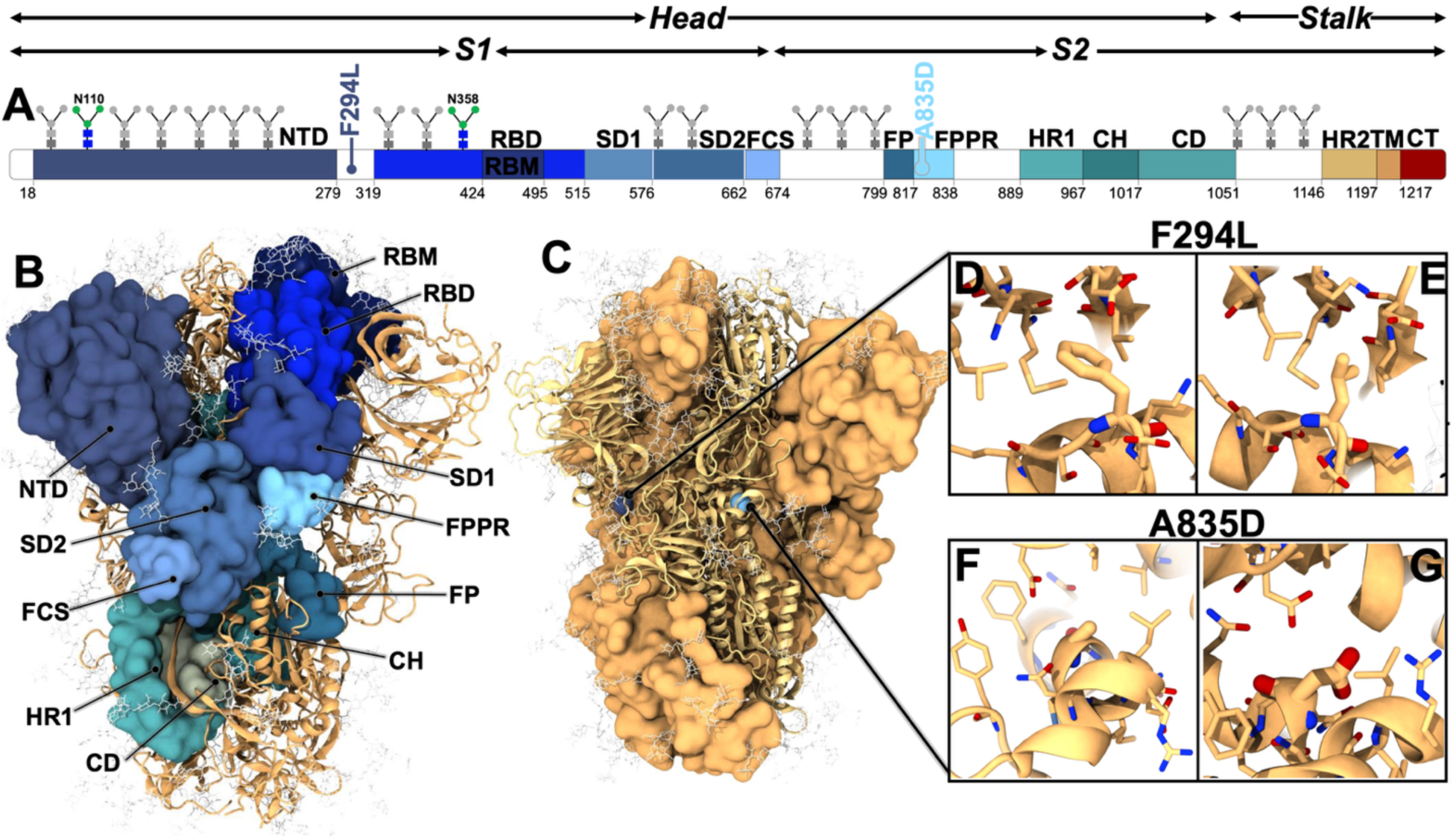
Modeling and system overview of the SHC014 spike GP. **(A)** Schematic of spike domains in a 1-dimensional sequence of the spike protein, including the F294L and A835D mutations, as well as glycans present at positions N110 and N358 (would be N370 in SARS-CoV-2), glycans not present on the human SARS-CoV-2 spike. Domains are defined as such: head (18 to 1124), stalk (1125 to 1256), S1 (18 to 668), S2 (669 to 1124), N-terminal domain (NTD, 18 to 279), receptor binding domain (RBD, 319 to 515), receptor binding motif (RBM 424 to 495), subdomain 1 (SD1, 515 to 576), subdomain 2 (SD2, 577 to 680), furin cleavage site (FCS, 663 to 674), fusion peptide (FP, 799 to 817), fusion peptide proximal region (FPPR, 818 to 838), heptad repeat 1 (HR1, 889 to 967), central helix (CH, 968 to 1017), connecting domain (CD, 1018 to 1051), heptad repeat 2 (HR2, 1146 to 1197), transmembrane domain (TM, 1197 to 1216), and cytoplasmic tail (CT, 1217 to 1256). **(B)** The 3-dimensional structure of the closed spike head highlights domains depicted in panel A. Glycans are shown in light gray licorice **(C).** Positions of F294L and A835D mutations are shown in blue and red spheres, respectively, on the spike head structure **(D, E, F, G)** Amino acid residues within 5Å of the F294L and A835D mutations.

Previous studies have demonstrated that mutations within the SARS-CoV-2 spike RBD, such as N501Y and T372A, impact infectivity and host-to-host transmission by modulating RBD conformations and ACE2 binding affinity^39–44^. Notably, a distal mutation far from the RBD, known as the D614G mutation (located ∼75 Å from the RBD’s center of mass in closed state), was associated with an increased infectivity and rapidly became the predominant circulating strain during early stages of the COVID-19 pandemic.^28,45–49^ Accumulating evidence predicts that the D614G mutation disrupts a key salt bridge to K854 within the neighboring chain’s S2 FPPR, thereby enhancing the transition of the RBD from its closed to up/open state.^28,50^

We have recently reported how a novel point mutation in the NTD (F294L) **(Figure 1C,1D,1E)** of the SHC014 spike protein increases viral infectivity.^51^ We further identified a second mutation in SHC014 spike’s FPPR region (A835D) (**Figure 1C,1F,1G**), which also enhances spike accessibility to ACE2 binding and increases viral entry into human host cells.^51^ However, the molecular mechanism by which these two mutations alter SHC014 viral infectivity in human cells remains incompletely understood, which is the central goal of this current work. Herein, we investigated the conformational plasticity of the glycosylated SHC014 spike ectodomain using multiple μs-long, conventional molecular dynamics (MD) simulations. Our atomistic simulations reveal how F294L and A835D mutations led to structural rearrangements to enhance RBD opening by altering allosteric communication pathways. Altogether, our work deciphered mechanistic insights into the dynamics of an SL-CoV spike GP, the SHC014 spike GP, which may be explored for understanding species spillover events and the development of effective pandemic-preparedness countermeasures.

## Results and Discussion

To predict how F294L and A835D mutations rescue spike:ACE2 binding and SHC014 viral infectivity in humans, we first constructed glycosylated models of SHC014 spike GP WT and mutant head domains. In total, our investigated models include the WT, two single mutants (F294L and A835D), and a double mutant (DM, F294L/A835D). To construct these models, we utilized our previously reported high-resolution Cryo-EM structure of the SHC014 spike GP head (PDB ID: 9CAS) domain in the closed state.^51^ While our reported structure was near complete in resolution, there were some missing loops (39 unresolved loops in the trimeric structure). Thus, to obtain a full head-domain WT construct, we employed a consensus approach to predict missing loop structures using AlphaFold^52^, SwissModel^53^ and Prime^54^, all of which were in good agreement (**Figure S1A-E**). Missing NTD loops (**Figure S1B**) were then grafted from the predicted Swiss Model, which again were in good agreement with AlphaFold and Prime models, to the Cryo-EM structure. Additionally, several SD2 unresolved loops were independently constructed using MODELLER^55^ through ChimeraX^56^ (**Figure S1C**). The complete spike head model was then fully N- and O-glycosylated following the SARS-CoV-2 spike glycosylation profile as previously outlined by Casalino et al. (2020)^31^ and comparing to SHC014 glycoanalytic data.^57^ Of importance, we modeled two glycans, N110 and N358, found on bat SHC014 spikes but not on SARS-CoV-2 spikes (**Figure 1A**). Position N358 corresponds to position N370 on SARS-CoV-2 spikes. Due to a threonine to alanine mutation at position 372, ablating a spike glycan sequon, the SARS-CoV-2 spike does not have a glycan at this position, whereas SARS-CoV spike does have a glycan at this N370 position.^42,43^ The two single mutants and the DM were generated from the WT construct using the psfgen mutate command in Visual Molecular Dynamics (VMD).^58^ Following the construction of the glycosylated SHC014 WT and mutant spike head domains, the models were explicitly solvated and neutralized (150 mM NaCl) with system sizes totaling ∼1 million atoms each **(See Table S1 for complete system information)**. We then conducted triplicate 1μs-long all-atom production MD simulations (following a total of 55 ns equilibration period) for each glycosylated ectodomain model **(Table S2)**, yielding an aggregate sampling time of 12 μs, and tracked the stability of the simulations by computing the root mean square deviations **(Figure S2)**. See **Table S3** for the complete SHC014 glycosylation profile and the Methods Section for complete model building and simulation details.

### Altered dynamics between SHC014 spike domains in WT and mutant strains

The Spike RBD conformational landscape, including RBD opening pathways, correlates with intra- and inter-protomer motions such as distances between the three RBDs, distances between RBDs and neighboring NTDs, and motions of other subdomains with the spike protein.^59^ To delineate structural movement within the SHC014 spike, we analyzed the root mean square fluctuation (RMSF) profile of the SHC014 spike protein across all the simulated models. We found that global RMSF profiles were broadly similar across all SHC014 spike variants (**Figure 2A**). However, we observed pronounced fluctuations within NTDs, RBDs, RBMs, SD2s, FCSs, and FPPRs.

**Figure 2:**
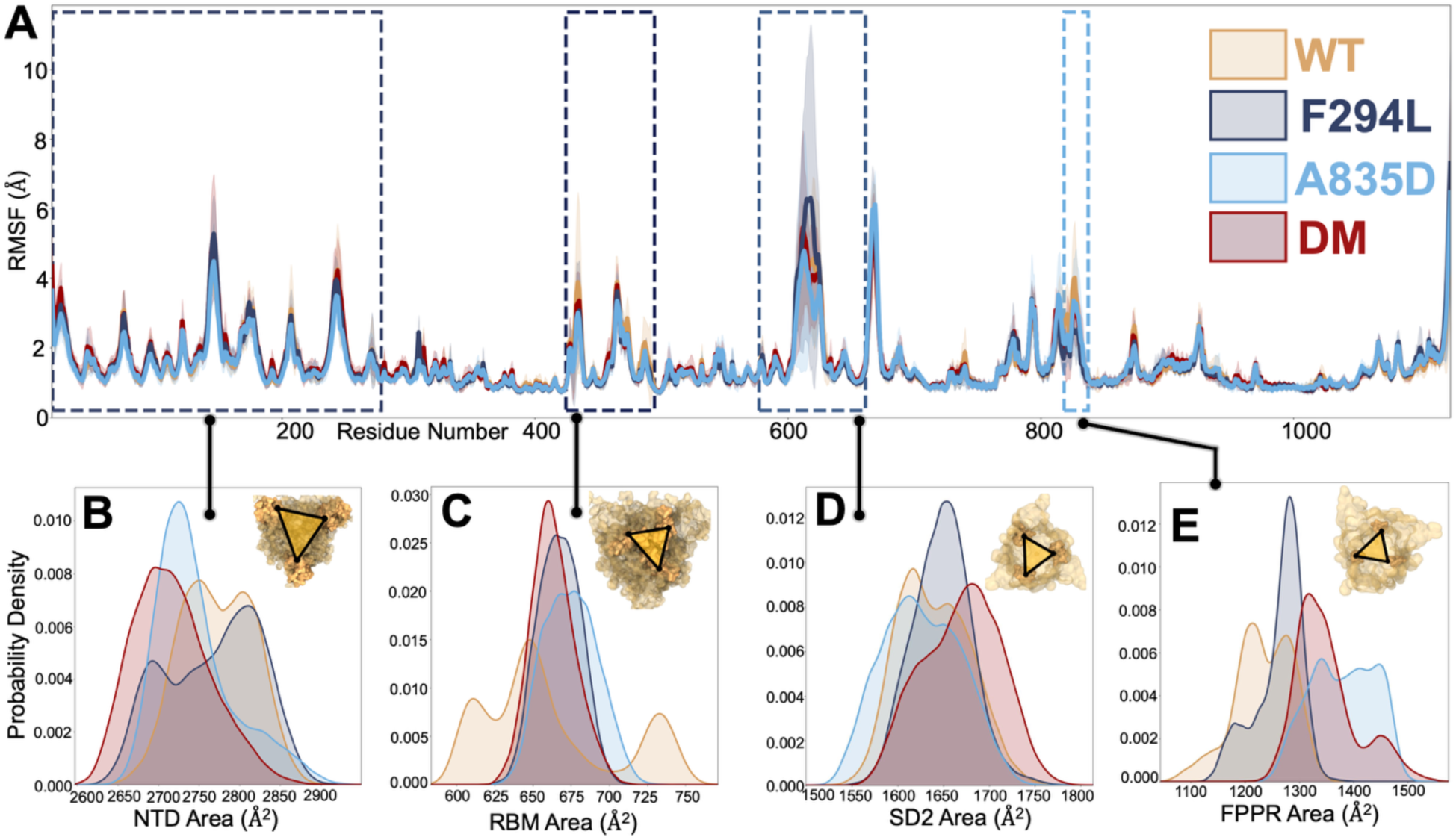
Characterization and flexibility of different domains of SHC014 spike FP. **(A)** Root mean square fluctuations of the entire protein system for each variant. **(B, C, D, E)** Distribution of areas in the domain with noticeable flexibility, such as the NTD, RBM, FPPR, and SD2. The variants are shown as WT in Gold, F294L in dark blue, A835D in cyan, and DM in red.

Based on the overall distribution of per-residue flexibilities (**Figure 2A**), we identified the domains with the most noticeable flexibility for more detailed dynamical analysis to unravel how their local motion influences global spike dynamics.

To investigate relative mobility of each domain of interest in the trimeric SHC014 spike, we identified the center of mass of each domain and tracked the area between each domain’s center of mass **(Figure S3)**. This metric indicates spatial breathing between each of the trimetric domains: a larger area indicating that domains are more widely spaced from one another, smaller areas indicating tighter packing towards the spike core. Tracking such areas over the course of all simulations allows us to determine the degree of breathing between domains as can be seen by the widths of these distributions. **Figure 2B** shows the distributions of areas between NTDs for the wildtype (WT) and the mutant spikes simulated in this work. WT SHC014 spike NTDs sample a broad and bimodal distribution of areas between them with a mean value of 2772A^2^ ± 45A^2^. F294L SHC014 spike NTDs also exhibit a broad, multimodal distribution of areas between them, with a mean value of 2767A^2^ ± 58A^2^. Contrarily, the A835D and DM constructs exhibited distributions with distinct skewing towards lower areas between their NTDs, with averages of 2746A^2^ ± 49A^2^, 2717A^2^ ± 48A^2^, respectively. Furthermore, the DM spike samples a distinctly tight distribution of NTD areas compared to WT, F294L, or the A835D spikes.

Similarly, the WT SHC014 spike RBMs exhibit a drastically different distribution of areas compared to A835D, F294L, and double mutant constructs (**Figure 2C**). WT RBMs exhibit a broad trimodal distribution of areas, with an average of 661A^2^ ± 43A^2^ with local maxima at ∼625A^2^, ∼∼670A^2^, and ∼750A^2^, indicating that it is traversing several conformational states, suggesting that WT SHC014 RBM (**Figure 2C**) and/or RBD (**Figure S4**) may not be readily accessible to ACE2 binding. However, all variant spikes indicate relatively narrow, unimodal distributions of RBM areas with averages of 2767A^2^ ± 58A^2^, 2745A^2^ ± 49A^2^, and 2717A^2^ ± 47A^2^ for the F294L, A835D, and DM spikes, respectively. Unimodal and narrow RBM (**Figure 2C**) and RBD (**Figure S4**) area distributions indicate ACE2 accessible conformations, suggesting increased availability of the RBDs for receptor binding relative to the WT. The F294L mutant exhibited a defined distribution with a tight peak, demonstrating exposure of its RBM, which is necessary to drive viral infectivity. Moreover, the A835D mutant displays a slightly larger RBM-area distribution compared to F294L, and the DM spike reveals the narrowest distribution with tighter packing (left shifted) compared even to the other variants, again indicating emergent interplay between A835D and F294L mutation positions. Given that the A835D mutation is in the FPPR and its distance to the RBD, these results suggest this mutation impacts RBD dynamics through allostery. Interestingly, the double mutant shows a sharp peak around lower RBM-areas, suggesting DM RBMs are held more compactly in this spike relative to other variants, making the DM RBMs much more accessible for receptor engagements, thereby driving cell entry. These differences in RBM area distributions, suggesting overall orientations to ACE2 binding, are consistent with our previous experimental findings as outlined by Tse et al., 2024^51^, where the degree of SHC014 infectivity is in the following order: DM > A835D > F294L > WT.

We also observe that subdomain-2 (SD2) exhibits variant-dependent differences in motion. In the SARS-CoV-2 spike, SD2 is a structural subdomain within the S1 region of the spike head and encompasses two elements previously implicated in membrane fusion: the FCS and the 630-loop. RMSF analysis (**Figure 2A)** shows that residues within SD2 display sequence-dependent flexibility, a result described in more detail in later sections. Accordingly, we also monitored the extent of SD2 “breathing” in the Bat CoV SHC014 spike for comparison. All SD2s demonstrate relatively broad distributions in trimeric areas between them (**Figure 2D**). Furthermore, the WT, A835D, and DM spikes demonstrate multimodal distributions in SD2 breathing with mean values of 1640A^2^ ± 39A^2^, 1626A^2^ ± 43A^2^, and 1670A^2^ ± 42A^2^, respectively. However, F294L exhibits narrower unimodal distribution centered around 1648A^2^ ± 32A^2^. In SARS-CoV-2 spikes, the FPPR has been reported to adopt altered conformations as a function of RBD openness and potentially serves to stabilize the RBD in the closed conformation,^27^ and thus, we also tracked areas between trimeric FPPRs over all simulations and per variant. All spike structures exhibited dramatically different degrees of FPPR breathing relative to one another. Firstly, all FPPR trimeric area distributions are multimodal, indicating multiple stable conformations (**Figure 2E**). Secondly, WT and F294L SHC014 spike FPPRs occupy trimeric areas between 1233A^2^ ± 54A^2^, and 1263A^2^ ± 44A^2^ respectively, while A835D and DM spike FPPRs occupy trimeric areas between 1380A^2^ ± 59A^2^ and 1351A^2^ ± 59A^2^, revealing little overlap between the variants with the A835D mutation (A835D and DM) versus the variants without the A835D mutation (WT and F294L). Additionally, F294L exhibits a clear primary conformation with a tight distribution of FPPR trimeric areas around 1263A^2^ ± 44A^2^ with two lesser populated shoulder distributions to the left of that major peak. Interestingly, DM spike FPPRs appear to adopt packing characteristics of both F294L, with a maximum population around 1351A^2^ ± 59A^2^ overlapping with F294L’s maximum, and A835D with the right-shift in ranges aligning to the A835D distribution. The A835D mutation is located within the FPPR, **Figure 1F,1G**; therefore, we expect local packing in this region to be altered due to this mutation. However, it is surprising to see the impact of F294L on FPPR packing towards the spike core and again indicative of the 294 position altering spike dynamics via allostery.

Collectively, these domain-based data reveal that the F294L and A835D mutations drive altered inter-domain dynamics. Particularly, we see WT spike RBMs are far more flexible in trimeric areas relative to variant spikes. Previously, we have reported that the WT SHC014 has low-to-no infectivity in human cells compared to the mutant strains.^51^ The WT RBM’s sampling multiple conformations compared to the other mutant strains could indicate that it is less accessible to ACE2, contributing to the observed decreased infectivity, as we hypothesized previously.^51^ Moreover, the observed altered dynamics of spike NTDs, SD2s, and FPPRs per spike variant indicate this sequence mutations could be impacting inherent spike allosteric networks.

### DM SHC014 Spike Demonstrates Increased Propensity for RBD Opening

The initial starting structure of the SHC014 spike glycoprotein (PDB ID: 9CAS) used herein was resolved with all three RBDs in the ‘down’ state, i.e., a “closed” spike conformation.^51^ Before the spike can bind to human ACE2, at least one of the RBDs needs to emerge from the down/shielded state into the up and open states to reveal the RBM.^30^ However, the SARS-CoV-2 spike’s RBM, and the RBD in general, is highly antigenic, eliciting the largest humoral immune response relative to any other domain within the spike protein.^60^ Thus, probabilistic tuning of spike RBMs in down versus up conformations has been hypothesized to provide replication advantage by balancing binding affinity and cell membrane fusion with immune recognition.^61^ Therefore, another factor of viral fitness and spillover to humans lies in the rate of RBD opening and resulting equilibrium distributions of down versus up RBDs. Thus, we sought to identify whether WT or variant SHC014 spikes could be observed to begin RBD opening within our μs-long conventional MD simulations. Experimentally determined RBD opening rates are reported to occur on a seconds-long timescale in vitro.^62^ Therefore, observation of spike RBD opening via conventional MD is likely intractable due to rare event sampling requirements, especially for such a large glycoprotein.^30^ However, we can use observations from conventional MD to inform on the tendency of spike RBDs towards opening. To track RBD position relative to that of an open RBD, we computed the distance between the center-of-mass of RBD β-sheet Cα atoms and the center-of-mass of CH’s C∝ atoms (a.k.a., the “spike core”), for each of three RBDs per spike and for every frame in all simulations. We herein refer to such values as “RBD-core distances” **(Figure S5)**, in keeping with our previous work,^30,31^ where larger and smaller values indicate increased or decreased displacement of the RBD from the spike core, respectively.

WT RBDs demonstrate a bimodal distribution with a primary, high probability conformation, and a distinct, low probability, left-hand tail (**Figure 3A**) where all the RBDs are tightly compacted towards the spike core in a “locked down” state. The F294L and A835D distributions overlap significantly with the RBD-core distances observed for WT RBD’s primary conformation. However, F294L exhibits a multimodal distribution of states within this range. The RBD-core distance of SARS-CoV-2 spike structure^17^ has been reported to be in the following range: closed state < 55Å, transition states between closed and open states 55 - 68Å, and the open state between 70 - 82Å.^30,63^ WT, F294L, and A835D all have an average RBD-core distance of 57Å ± 1Å, while that of DM is 58Å ± 1Å. Interestingly, the DM spike RBD samples a population of conformations with a slightly increased RBD-core distance, potentially exhibiting a slightly heightened propensity for opening, which is crucial for SHC014 infectivity in humans (**Figure 3A,S5**). The DM “initiating up” state exposes the spike to neutralizing antibodies (**Figure 3B,3C,3D**). We have previously shown that the SHC014 DM spike demonstrated an increased sensitivity to neutralization by an antibody called Adagio-2 (ADG2)^51^, hypothesized to be due to an increased population of open RBD conformations. To further understand how accessible the RBD is to neutralizing antibodies and the glycan shielding effect, we conducted accessible surface area (ASA) calculations for RBM and other domains with a probe radius of 7.2Å. The ASA results indicate that there are only subtle differences in the ASA of the given domains (**Figure S6).** Altogether, we posit that the F294L and A835D mutations confer a fitness trade-off in SHC014-CoV by increasing RBD opening to facilitate cell entry while simultaneously enhancing susceptibility to neutralizing antibodies.

**Figure 3:**
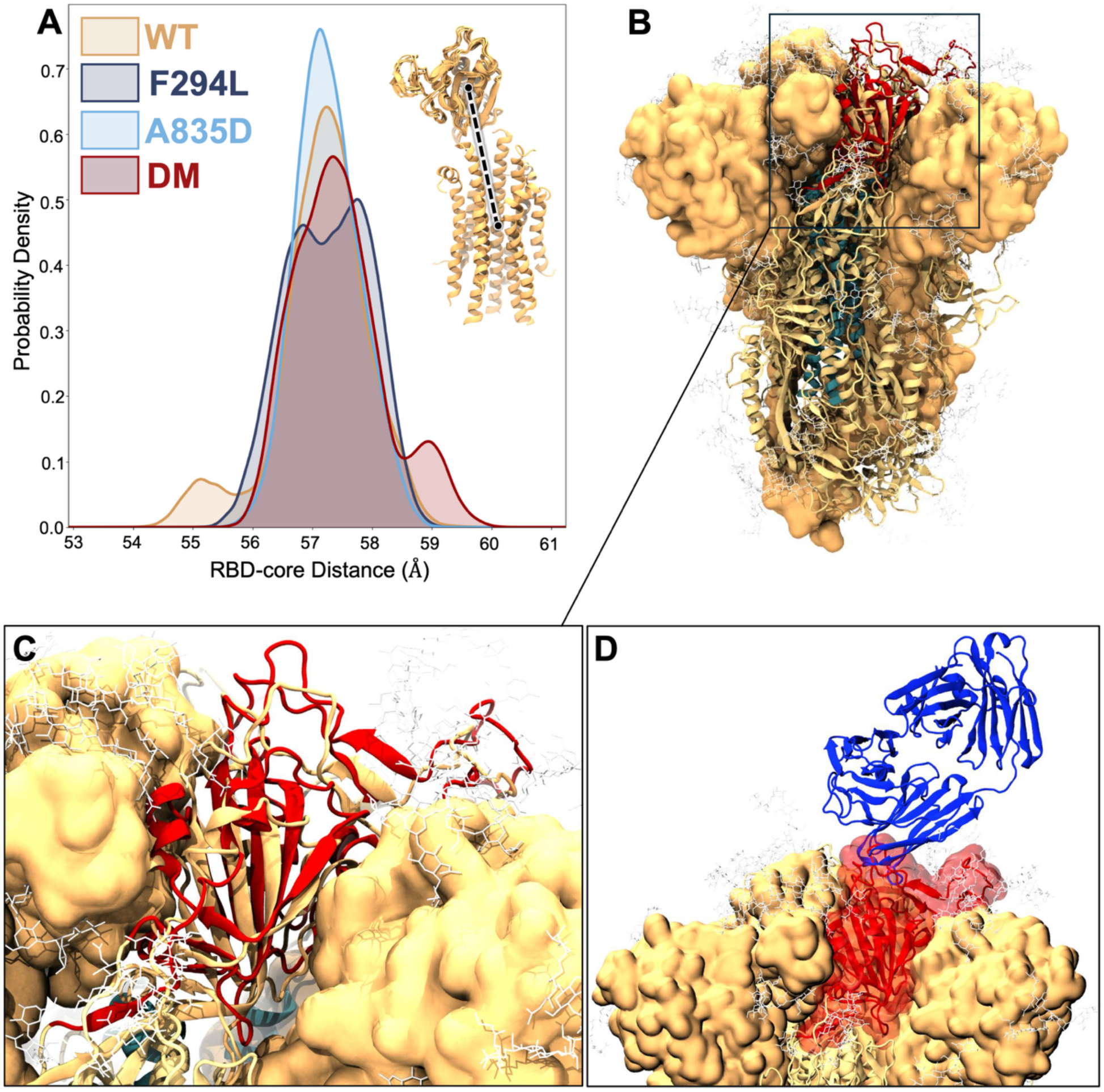
DM spike RBD initiates opening in conventional MD simulations. **(A).** RBD-core distance distribution plot, which was calculated based on the center of mass of RBD to the central helices of the spike core. **(B, C)** A close view of a set of distributions in the DM spike RBD that demonstrated a slight increase in RBD opening; WT is shown in gold and DM in red **(D).** Complex of SARS-CoV-2 neutralizing antibody (ADG2) in complex with the DM spike RBD. ADG2 antibody is shown in blue.

### Mutant spike contact networks drive altered domain dynamics

We next sought to characterize the atomic-scale interactions impacted by the F294L and A835D mutations and predict how those contacts may impact spike domain motions, including RBD opening. Previous studies have shown that as the SARS-CoV-2 spike RBD transitions between down, up, and open conformations, several salt-bridging and hydrogen bonding interactions form and break,^30^ including within the FPPR.^63^ As such, we identified key contacts between positions 294 and 835 and their neighboring amino acids and glycans, and we tracked such contacts (heavy atoms within 4Å of one another) over the course of all simulations and for all studied spike variants (**Figure 4,S7-S15)**.

**Figure 4:**
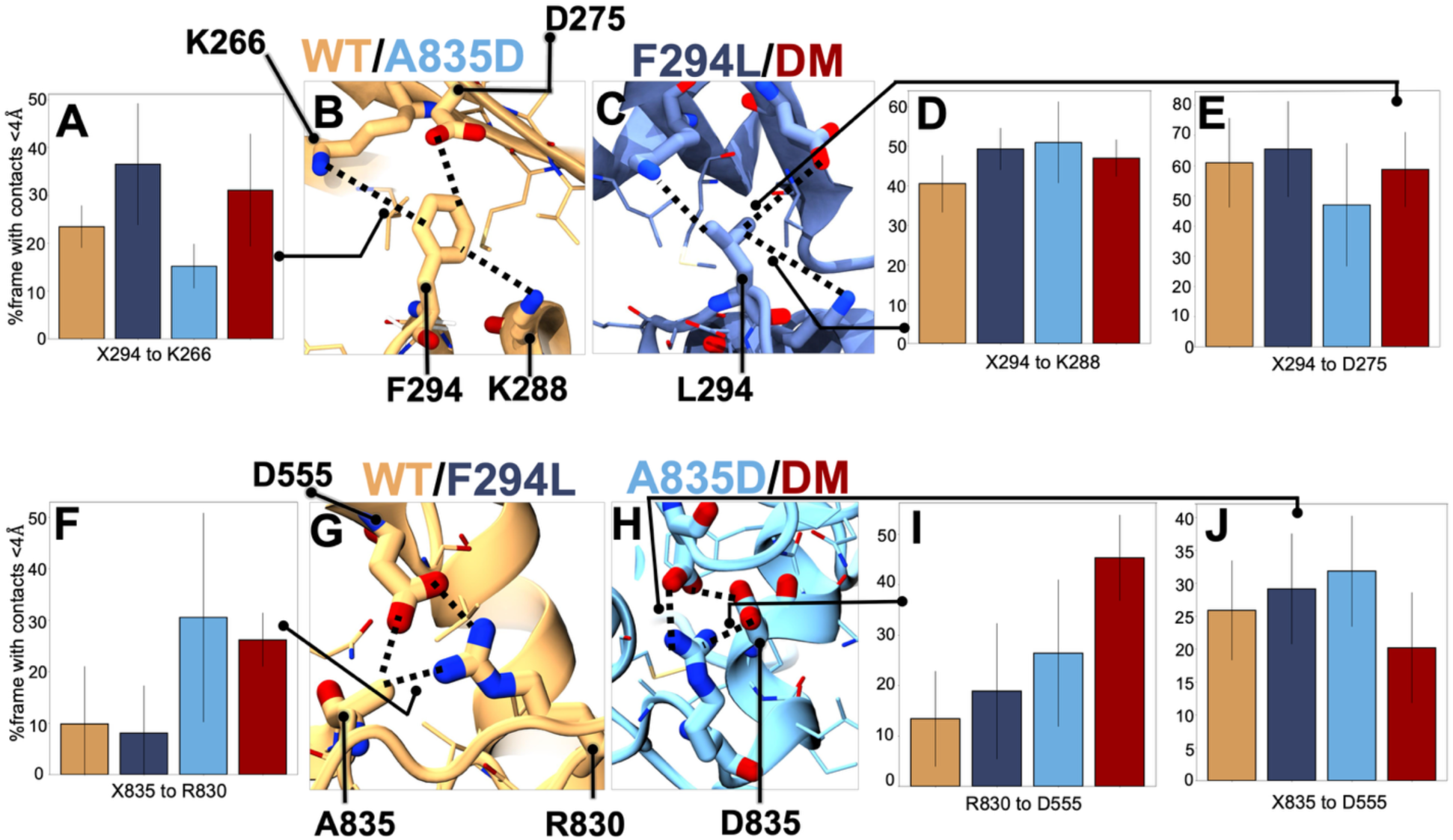
Contact analysis of residues within 4Å of (B, C) F294L and **(G, H)** A835D mutations. Contact frequency less than 4Å between **(A)** X294 to K266, **(D)** X294 to K288, and **(E)** X294 to D275, where X at position 294 can be phenylalanine or leucine. **Figures F, I, and J,** represent contact frequency between X835 to R830, R830 to D555, and X835 to D555, respectively where X can be Alanine or Aspartate.

Around position 294, there are several positively charged Lys and negatively charged Asp residues **(Figure 4B,4C)**. Because aromatic amino acids in this region could exploit cation-π or anion-π interactions for stability, we sought to track contacts between these charged residues and position 294 **(Figures 4A,4D,4E,S7-S9)**. Firstly, we see that contact between L294 and K266 is ∼10-15% more favorable than F294 to K266 (**Figure 4A,S7)**. Secondly, we see that contacts between X294 and K288 are largely the same regardless of F/L at this position (**Figure 4D,S8)**. Thirdly, we see the A835D variant has a slightly decreased propensity for contact between F294 to D275, relative to the other variants (**Figure 4E,S9)**.

There are several Asp and Arg residues in the local region surrounding position 835, and we also tracked contacts between these neighboring residues (**Figure 4G–4J,S10-S12)**. We predicted in previous work that the introduction of another Asp in the FPPR region may therefore result in marked rearrangement within the area to alleviate electrostatic clashing.^51^ Our simulations have supported this hypothesis: D835 is very close to the neighboring chain’s D555, which sits at the base of the SD1, close to the RBD (**Figure 1B**). We observed a ∼20% increase in contact frequency between D835 and R830 relative to A835 to R830 contacts seen in WT and F294L variants (**Figure 4F,S10)**. Moreover, we see ∼32%, ∼26%, and ∼18% increase in R830 to D555 contact frequency for the DM spike relative to WT, F294L, and A835D spikes, respectively (**Figure 4I,S11)**. Interestingly, we see that although X835 and D555 are in close proximity in all simulations -- ∼25%, ∼28%, ∼32% contact frequency between X835 and D555 in WT, F294L, and A835D, respectively – there is a distinct decrease in X835 and D555 contact frequency for DM spikes to ∼19% (**Figure 4J,S12)**. To unravel the formation and persistence of the salt-bridge triad between R830-D555-D835, we computed the area between the center-of-mass of each residue throughout all simulations and for contacts spanning each of the protomeric interfaces. From these results, we see that 38% and 44% of simulation frames from A835D and DM spike variants, respectively, show tight salt bridge triad formation (area < 30 A^2^), which is absent in WT and F294L variants due to the presence of Alanine at position 835 **(Figure S13).** These data suggest two potential mechanisms for alleviation of the densely packed negative charge: (1) recruitment of R830 to stabilize D835 and D555 as they neighbor each other, and (2) adjustment of RBD position to separate D835 and D555, as this was the only spike seen to begin initiation RBD opening.

Additionally, we observed protein-glycan contacts, particularly between R830, which mediates the salt bridge interactions, and its neighboring glycans N270 and N590, with differences across the variants **(Figure S14-S15)**. Taken together, we posit that the A835D mutation introduces dense packing of negative charge in the immediate area surrounding the FPPR and SD1/base of the RBD. To alleviate this negative charge, A835D and DM spikes recruit R830 to this region for better charge balance. We believe the F294L mutation contributes to a loosening of interactions at the base of the NTD, and particularly at the base of a loop, known as the N2R linker, which is responsible for passing allosteric information between the NTD and RBD. We believe the DM spike is therefore marginally more adept at alleviating negative charge density via opening its RBD, thus contributing to the slight increase in proportion of DM RBDs initiating transition to the up state.

### 620-loop Exhibited Enhanced Dynamics in Double Mutant System

Dynamics and positioning of the “630-loop” in SARS-CoV-2 spike (residues 620 to 640) have been shown to correlate significantly with RBD up/down conformational states.^64^ Moreover, perturbations in or near this loop, including the D614G mutation, to alter its flexibility/compactness have been shown to directly impact RBD conformations.^28,65^ Based on these insights from the SARS-CoV-2 spike, we hypothesized that the structurally analogous region in the SHC014 spike, which we have called the “620-loop”, may similarly modulate allosteric communication between the NTD, RBD, and FPPR. Per-residue flexibilities for the 620-loop **(Figure 5A,5B)** as a function of spike variant indicate that while all spike 620-loop exhibit significant degrees of flexibility relative to other spike domains, the F294L variant demonstrates remarkable variance in flexibility in this region, **Figure 5C**.

**Figure 5:**
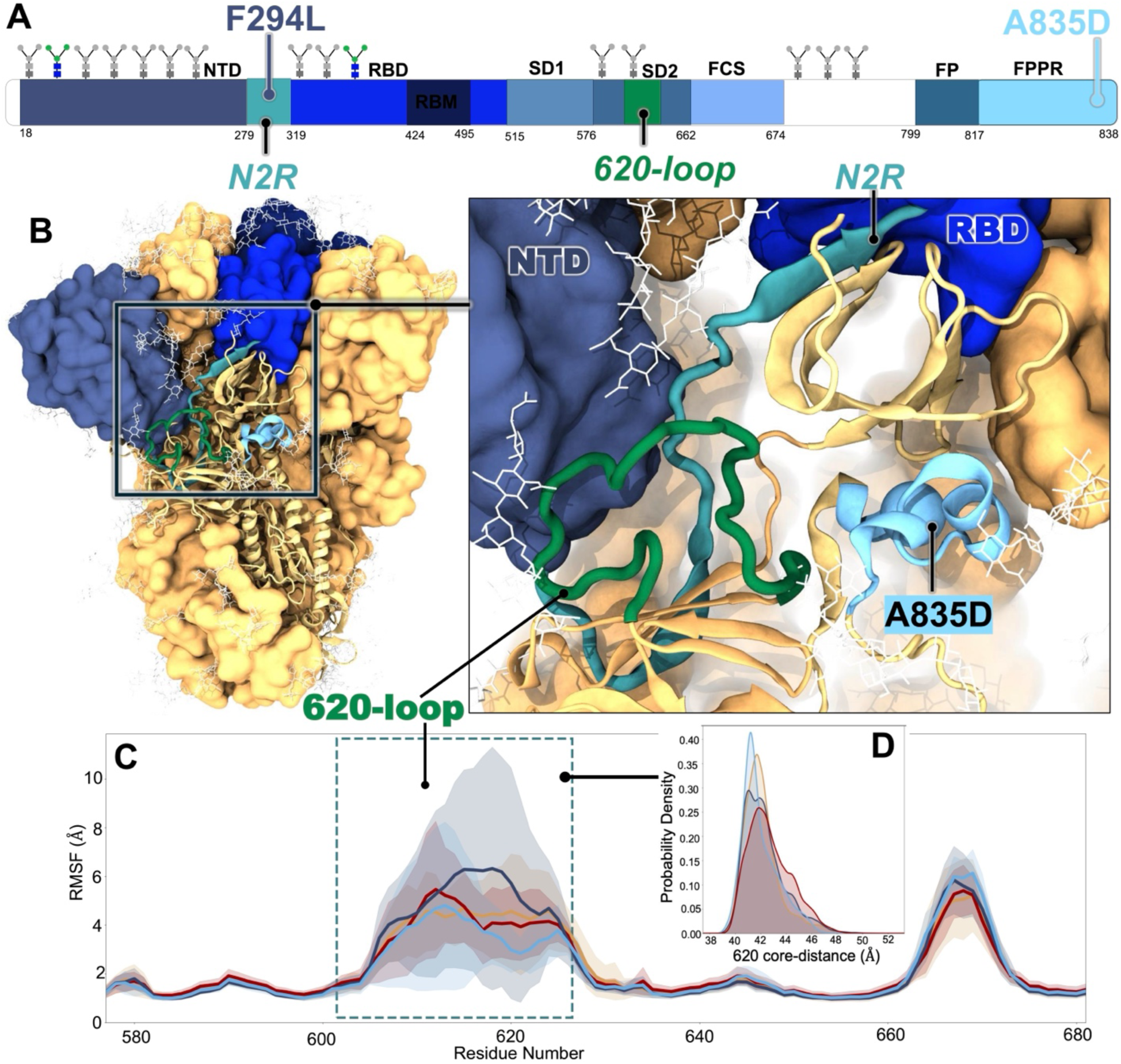
The 620-loop exhibited varying degree of flexibility. **(A).** 1D sequence of SHC014 spike protein from the NTD to the FPPR region only. The 620-loop is shown in green. **(B)** Close view of the 620-loop and the N2R linker **(C)** RMSF plot depicting per-residue flexibility within the SD2 domain with a particular focus on the 620-loop. **(D)** 620-core distance plot, which was calculated based on the center of mass of 620-loop to the central helices of the spike core.

To further characterize the behavior of the 620-loop in SHC014, we tracked how much the 620-loop deviates from the center of the SHC014 spike protein core. To do so, analogous to the RBD-core distances, we calculated the 620-core distances as the distance between the 620-loop Cα atoms’ center-of-mass and the CH Cα atoms center-of-mass. 620-core distances from all SHC014 spikes occupy right-skewed probability distributions with similar averages of 42.0Å ± 1.45Å, 42.31Å ± 1.63Å, 41.96Å ± 1.43Å, and 42.72Å ± 1.69Å for WT, F294L, A835D, and DM spikes, respectively **(Figure 5D)**. However, DM spike appears to sample larger 620-core distances more frequently than the other spikes, which may correlate with the initiating up-RBD conformations observed for this variant **(Figure S16A-S16B)**. The Y623H mutation, positioned within the 620-loop of sub-domain 2 in SHC014 GP, has been previously shown to enhance viral entry, potentially driving the opening of the RBD^66^. While preparing this manuscript, Tse et al. (2025)^67^ reported that the Y623H substitution may modulate the propensity of SHC014 RBD to sample an open conformation, thereby increasing viral infectivity by enhancing spike:ACE2 binding. Altogether, our work complements these findings by uncovering the varying degree of flexibility in the 620-loop across the variants, which may allosterically alter RBD conformations **(Figure S16C-S16D).**

### A835D mutation reshapes long-range allosteric communication

To understand how the F294L and the A835D mutations reshape allostery within the SHC014 spike, particularly in altering domain-domain communication critical for RBD opening, we carried out dynamical network analysis using the Weighted Implementation of Suboptimal Paths (WISP) method^68^. Using a weighted residue network, we represented each residue as a node and connected pairs of residues (edges) with weights proportional to the strength of their correlated motions. This graph-based method not only uncovers how a group of nodes that form a domain can establish networks with distant regions within the spike but also how the two-point mutations studied herein can influence patterns in allosteric communications and structural rearrangements **(Figure 6A,6B)**. From our dynamical network analyses, we extracted edge-weighted domain-domain pairs **(Figure S17)** to map a continuous allosteric pathway, and we noted differences in communication routes across the variants. Interestingly, SD1, positioned at the base of the RBD **(Figure 6B)**, serves as a critical allosteric hub where nearly all identified communication pathways converge, thereby prompting it to act like a ‘molecular junction’ that relays information between functionally distinct regions of the spike. Just like a traffic junction that allows multiple roads to intersect, SD1 relays information from either the NTD or the SD2 to the RBD. Previous structural studies of the full-length SARS-CoV-2 spike suggested SD1 to be a structural relay for communication between RBD and FPPR, and capable of sensing displacement on either side.^27^ Similarly, computational modeling of the spike protein using coarse-grained simulations has reported SD1 as an allosteric center between RBD and FPPR.^69^ Therefore, our findings corroborate the existing body of evidence that SD1 acts as an important allosteric region.

**Figure 6:**
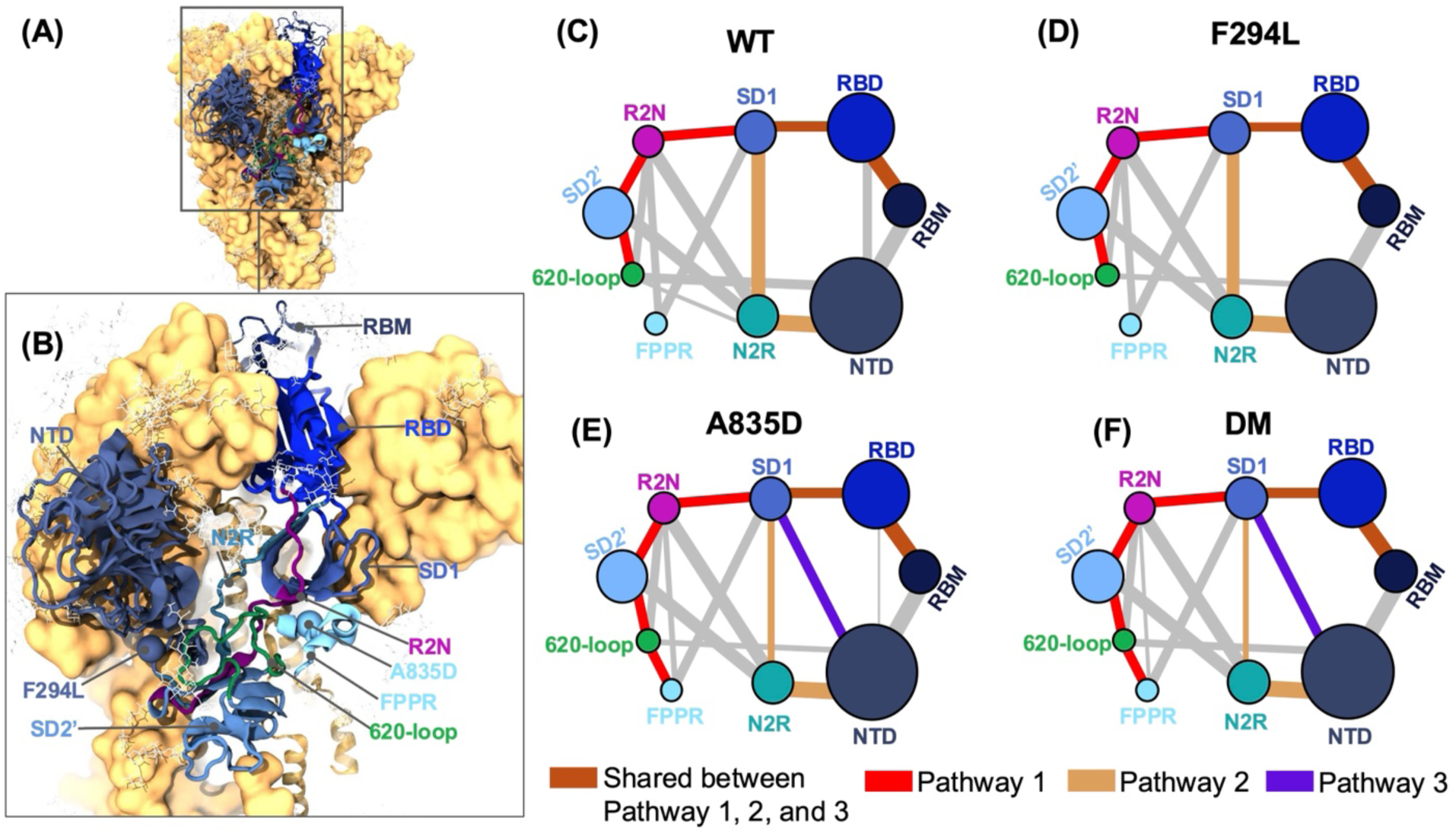
Dynamical network analysis reveals communication pathways across the SHC014 spike variants. **(A)** Domains within the allosteric routes and **(B)** a close view of the domains, including the N2R and R2N linkers. **(C, D, E, F)** A residue-based graph network depicting how domains communicate with each other, weighted by their edges for WT, F294L, A835D, and DM, respectively. **(D)** Each node represents domain size, and edge thickness corresponds to communication strength. Brown edges highlight the shared route between all three pathways, red represents pathway 1 in WT and F294L variants in the following order: 620-loop– SD2’ – R2N – SD1 – RBD – RBM as well as A835D and DM which extend the communication in the following order: FPPR – 620-loop– SD2’ – R2N – SD1 – RBD – RBM among the variants. Orange represents allosteric communication along Pathway 2 in the following order: NTD – N2R – SD1 – RBD – RBM, while the purple depicts Pathway 3 seen only in A835D and DM: NTD – SD1 – RBD – RBM. The gray edges indicate non-primary domain-domain pair communications pathways that may be utilized by SHC014 spike protein during allosteric communications. SD2’ denotes the SD2 domain without the 620-loop.

The R2N linker, which connects the RBD to the base of the NTD and SD2, has been shown to primarily facilitate allosteric communication in the Delta and Omicron SARS-CoV-2 spike variants.^63^ Notably, both A835D and DM displayed a direct allosteric pathway from the FPPR via the SD2 to the R2N linker, then the SD1, which acts as the allosteric bridge, ultimately relaying the information to the RBD **(Figure 6E,6F)**. The SD2 contains both the FCS and the 620-loop; therefore, to understand the role of the 620-loop in the allosteric networks, we selected both the 620-loop alone and the SD2 with the FCS, hereafter referred to as SD2’. We identified three main communication pathways to the RBD named “Pathway 1”, “Pathway 2”, and “Pathway 3”. In the WT and F294L variant, Pathway 1 shows baseline allosteric communication from the 620-loop to the RBD **(Figure 6C,6D)**. Notably, in the A835D and DM variants, pathway 1 starts at the FPPR, then proceeds to the 620-loop, and ultimately to the RBD, indicating how the D835 mutation extends the baseline pathway, enabling FPPR to contribute to allosteric communication with the RBD **(Figure 6E,6F)**. Similarly, the D614G mutation, located far from the RBD, has been reported to reshape allosteric networks in the SARS-CoV-2 spike.^63^ Moreover, the FPPR to 620-loop to RBD crosstalk observed in A835D and DM Pathway 1 is consistent with earlier findings **(Figure 5D, S16A,S16B)** showing enhanced flexibility in the DM 620-loop compared to other variants. These results delineate how point mutations distant from the RBD, such as the A835D with the FPPR, can modulate both structural dynamics and long-range allosteric communication in the spike.

Furthermore, the N2R linker has been proposed to facilitate communication in the SARS-CoV-2 spike S1, leading to RBD opening^59,70^. In all SHC014 spike variants considered here, we also identified a conserved communication route, hereafter referred to as Pathway 2, running from NTD-N2R-SD1-RBD-RBM **(Figure 6C-6F)**. Moreover, given increased edge weights for SD1 to N2R nodes, Pathway 2 is more prominent in the WT and F294L compared to other variants, indicating that information flowing from the NTD to the SD1 has to first pass through the N2R linker. However, we observed that in A835D and DM, a “shortcut” is established hereafter referred to as “Pathway 3” **(Figure 6E,6F),** where information can flow directly from the NTD to the SD1, which rebalances the communication within the three edges instead of the two in WT and F294L. Kearns et al.^63^ reported distinct allosteric communication across human SARS-CoV-2 spike variants, with Delta and Ancestral spikes primarily transmitting information between the NTD and RBD via the R2N and N2R linkers, respectively, whereas Omicron was capable of utilizing both. Similarly, SHC014 DM and A835D variants may be capable of using both Pathway 1, 2, and 3 to potentially enhance RBD opening dynamics and increase ACE2 binding efficiency, as we have seen in previous experimental work.^51^

## Conclusions

In this work, we present multiple 1-μs-long explicitly solvated all-atom MD simulations of the fully glycosylated SHC014-CoV WT and variant spike ectodomains. Our previous work demonstrated that the acquisition of two novel mutations, F294L and A835D in the NTD and FPPR, respectively, enhances viral infectivity by increasing RBD availability to the host cell receptor, ACE2.^51^ Herein, we have observed that these mutations lead to the structural rearrangement of, and altered communication between, domains affecting the global motion of the spike protein. The WT RBM was found to sample multiple conformations, thereby making it less accessible to receptor binding compared to the DM, likely contributing to the WT’s low infectivity in an ACE2-dependent manner as observed experimentally.^51^ The structures of SARS-like bat coronaviruses, including SHC014, have been resolved with the RBD in “down” conformation.^51,66^ Notably, we also observed that the DM RBD initiates marginal opening even in conventional MD simulations, which is unusual given the timescales of spike opening (milliseconds), which typically requires enhanced sampling methods to observe.^30,62,63,71–75^ While the F294L and A835D mutations enhance ACE2 binding and cell entry, they also expose SHC014-CoV spike to neutralizing antibodies like ADG2, indicating a fitness trade-off and potential therapeutic avenue that may be exploited for the design of effective countermeasures to support pandemic preparedness efforts.

We observed that initiation of the RBD opening event in the DM may be allosterically modulated by the increased flexibility of the 620-loop and its crosstalk. Consistent with our findings, previous studies have shown that the Y623H mutation^66,67^ within the 620-loop of SHC014-CoV led to an increase in viral infectivity compared to the WT, further affirming the role of this loop in RBD openness to enhance cell entry. The 620-loop in SHC014-CoV corresponds to the 630-loop in SARS-CoV-2 spike, which has similarly been shown to correlate with RBD motion.^63^ At the atomic scale, the F294L mutation appears to disrupt the network of π-mediated interactions, thereby enhancing the flexibility of the N2R linker and contributing to marked accessibility of RBD to ACE2. Contrarily, we found that the DM alleviates the dense negative packing around the FPPR region introduced by the A835D mutation to favor RBD transitioning to the open state. Through dynamical network analyses, we revealed three allosteric pathways via the R2N and N2R linkers as well as the NTD, all converging at the SD1 domain, which acts as an allosteric hub relaying information to the RBD. While Pathway 1 was present across all variants, it extended to include the FPPR region in the A835D and DM variants, suggesting that this pathway is strengthened in these mutants compared to the WT and F294L. This finding illustrates how distant mutations, such as A835D in the FPPR, can reshape long-range allosteric communication within the spike. In contrast, Pathway 2 requires information to pass through the N2R linker from the NTD to SD1. Although Pathway 2 was observed in A835D and DM, it exhibited reduced communication than in WT and F294L. Interestingly, a new route, termed Pathway 3, emerged in A835D and DM, establishing a direct shortcut from the NTD to SD1 that may balance the communication in Pathway 2. Altogether, our work provides atomistic insights into the structural dynamics and molecular adaptations governing WT and mutant SHC014-CoVs and probable therapeutic opportunities against SHC014 and related SARS-like bat coronaviruses for pandemic preparedness efforts.

## Materials and Methods

### Model system construction

The spike protein of SHC014 is a large, glycosylated trimeric structure with its ectodomain spanning residues 18 to 1124, which comprises the S1 and S2 subunits (**Figure 1A**, main text).^14,15^ We have previously reported the experimental structure of SHC014 spike protein (PDB ID: 9CAS) as described in Tse et al., (2024)^51^, with all three receptor binding domains in closed conformations used herein to construct the ectodomain, fully glycosylated models of the wildtype (WT) and three mutant spikes. Specifically, four models were constructed, including the WT, two single mutants -- the first mutation is in the N-terminal region (NTD) where phenylalanine at position 294 was mutated to leucine (F294L) while the second mutation is in the fusion peptide proximal region (FPPR) where alanine was mutated to aspartate (A835D) -- and a double mutant (F294L+A835D). Thus, we employed a multi-step modeling approach to construct a complete model of the SHC014 spike head domain with fully resolved loops, N- and O-linked glycans, and reversion of stabilizing mutations., *<u>Missing loops modeling:</u>* The resolved cryoEM structure of SHC014 GP contains several unresolved sequence gaps corresponding to flexible loops in the NTD, SD2, and FPPR, resulting in 39 unresolved loops in the trimeric structure (**Figure S1A**). To generate a complete ectodomain WT construct, missing gaps were modeled as disordered loops using various protein structure prediction tools, including the Swiss Model^53^, AlphaFold2^52^, Prime^54^, and MODELLER^55^. The NTD (residues 18 to 279) contains several unresolved flexible loops (9 per monomer, 27 in trimeric structure). Considering the poor density of these unresolved loops, we elected to employ Swiss Modeling, AlphaFold2, and Prime Homology Modeling and achieved consensus amongst the modeling techniques for these NTDs. Missing loops within the NTD were then grafted from the Swiss Model prediction – which was, again, highly similar to Prime and AF2 predicted models -- onto the Cryo-EM structure in each chain A, B, and C (**Figure S1B**). To model the SD2’s unresolved loops, per monomer, 5 complete SD2 models were independently generated using MODELLER, from which the top per-chain SD2 model was chosen based on z-score and taking care to avoid overlap with other domains. Resultant refined loops within each top predicted SD2 model were visually inspected and grafted to the resolved CryoEM structure (**Figure S1C**). To achieve the most accurate model of the FPPR, we identified the unresolved sections of the FPPR sequence from Uniprot (residues 818 to 831) and predicted the structure of this loop via homology modeling with Swiss Model using the human SARS-CoV-2 spike glycoprotein as a template structure (PDB 7JJI) for the SHC014 sequence in this region^76^ (**Figure S1D**). The complete FPPR predicted region was then, as done with the NTD, grafted onto the CryoEM resolved regions of the SHC014 structure. In all cases of grafting, the joined positions between grafted predicted loops and resolved CryoEM were visually inspected to ensure there were no clashes, overlaps, or ring penetrations. Notably, the S1/S2 FCS was modeled in an uncleaved state. Uniprot and comparison to human SARS-CoV-2 spike system structure were used to identify necessary disulfide bonds (15 per monomer) as listed in **Table S1**.

### Glycosylation/ Protonation/ Solvation/ Neutralization

SHC014 spike protein ectodomain glycosites were predicted using the NetNGlyc webserver^77^ and the N-Glycosite tool^78^ and further confirmed by previously reported SHC014 site-specific glycan analysis^57^ and comprise 18 N-glycan sequons (N-X-S/T) per monomer, including N110 and N358, but not in SARS-CoV-2. N110 and N358 correspond to N110 and N370, respectively in the SARS-CoV-2 glycosylation profile. Our modeled constructs are fully N-/O-glycosylated following a similar human glycoprofile as used by (Casalino et al., 2020)^31^, which was consistent with (Watanabe et al., 2020).^25^ Detailed glycosylation profiles of SHC014 GP per chain simulated herein are shown in **Table S3.** In summary, 57 glycans (18 x 3monomers N-glycans and 1 x 3monomers O-glycans) were incorporated into the WT spike system. Any protein/glycan clashes or ring penetration in each construct were remedied using VMD’s manual structure manipulation tool, Molefacture, and VMD’s site-specific minimization and equilibration MD simulation tool, AutoIMD.^79^

### System Preparation

It is important to note that in this study, the WT S1/S2 site was modeled in an uncleaved form. Consistent with (Casalino et al., 2020)^31^, the S1 and S2 subunits within each protomer have been assigned to the same chain (referred to as A/B/C). Glycans were assigned segnames from G1 to G57, as reported in **Table S3**. Protonation states for all titratable residues (Asp, Glu, Tyr, Lys, Cys, and Arg) were assigned using PROPKA^80^ at pH 7.4 and His states were assigned with Schrodinger’s Protein Preparation Wizard tool^81^. All Asp and Glu were determined to be deprotonated and negatively charged, all Cys were determined to be involved in disulfide bonds and thus not titratable, all Tyr were determined to be protonated and neutral, all Lys and Arg were determined to be protonated and positively charged. His numbered 38, 54, 59, 492, 662, 1041, and 1066 were modeled as neutral and singly protonated on the δ-Νitrogen (HSD). His numbered 196, 612, 642, 1031, and 1042 were modeled as neutral and singly protonated on the ε-Νitrogen (HSE). Protein structure files for the WT protein structure was generated via VMD’s psfgen module according to CHARMM36 all-atom additive force fields for proteins and glycans^82,83^. The mutate command in VMD’s psfgen module^58^ was then used to modify the WT structure to F294L, A835D, and DM spike structures. The WT construct was fully solvated with explicit water molecules described using the TIP3P model^84^ and neutralized with a 150 mM concentration of Na and Cl ions. See **Table S1** for complete details of atom counts and system sizes.

### Molecular dynamics (MD) simulations

The solvated and neutralized WT, F294L, A835D, and DM ectodomain models were minimized and briefly equilibrated using NAMD3^85^ with the protocol as follows. For all the following described protocols, system temperature and pressure were controlled with a Langevin thermostat^86^ and a Langevin barostat^87^, respectively. Periodic boundary conditions were used to model infinite bulk conditions, and particle-mesh Ewald^88^ was used to handle long-range electrostatic interactions. The SHAKE algorithm^89^ was employed to fix covalent bonds involving all hydrogen atoms. Van der Waals interactions and electrostatic interactions were computed with the following cutoff scheme: cutoff distance of 12Å, pair list generated for all pairs within 13.5 Å, and switching function implemented for all pairs between 10Å and 12 Å apart. Non-bonded interactions were excluded for bonded atoms within 1-4 bonds of each other. ***<u>Minimization and heating:</u>*** First, waters and ions were subjected to 10080 steps of conjugate gradient minimization, protein and glycan atoms were held fixed by Lagrangian multiplier constraints. Next, a molecular dynamics (MD) heating protocol was used to gradually introduce the systems to biologically relevant temperatures: protein and glycan atoms were held fixed during this step (Lagrangian multiplier constraints), starting at 10K, every 10080 MD steps (1fs/step) the temperature was incremented by 25 K until 310K. At 310K, 1ns of short equilibration MD was performed. ***<u>Volume and pressure equilibration:</u>*** Following heating protocol, Lagrangian constraints were removed from protein and glycan atoms and a short (2520 steps) conjugate gradient minimization was performed wherein all water and ion atoms were allowed to freely move and protein and glycan atoms were lightly restrained with a force constant k= 1kcal/mol/Å^2^. Following this short minimization, 0.5 ns (2 fs/step) of pressure and volume equilibration were performed wherein the periodic box was allowed to fluctuate in size, but protein and glycan atoms were still subjected to the light restraint (k=1kcal/mol/Å2). *<u>Free NVT equilibration:</u>* Following pressure and volume equilibration, the periodic box was held fixed (useFlexibleCell set to no, useConstantArea set to no) for ∼1 ns (2 fs/step) of free equilibration in which all restraints and constraints were removed from protein and glycan atoms. See **Table S2** for total simulation time.

### Analysis methods

#### Protein Root Mean Square Fluctuations

The Root mean square fluctuations for the wildtype and mutant SHC014 spike GPs were calculated using MDAnalysis^90,91^ RMSF module. Each trajectory was aligned onto the initial coordinates using the C_α_ atoms of the entire protein to remove global translation and rotational degrees of freedom. We further calculated the RMSF of the glycans by aligning the glycosylated trajectory to the reference frame using the heavy atoms of the protein glycosite (i.e., Asn’s for N-linked glycans and Ser or Thr for O-glycans). After which we calculated the RMSF for all C1 atoms in the glycan segment analogous to Cαs in protein segments and averaged those C1 RMSFs per frame over the whole simulation.

#### Residue to Residue Contact Distance Measurements

Contact analysis was carried out for every frame in each simulation across all chains of each variant using the distance module MDAnalysis.^90,91^ The minimum distance between heavy atoms was used to calculate distances for the following: X835 to D555, X835 to R830, R830 to D555, X294 to D275, X294 to K266, X294 to K288, R830 to N590 glycan, and R830 to N270 glycan. Only side-chain-to-side-chain contacts were considered for contact analysis involving X294 and its neighboring residues, as backbone atoms are already very close to one another.

#### RBD-core distances

Per-frame distances between the RBD in SHC014 spike and the spike core were computed using MDAnalysis^90,91^ by measuring the distance between the center of mass of the RBD Cα atoms (residues 319 to 515) and that of the spike’s central helices (Cα atoms from residues 968 to 1017). These calculations were performed per replica trajectories of each variant (WT, F294L, A835D, and DM).

#### 620-core distances

As described above, we carried out similar analysis between the center of mass of the 620-loop’s Cα atoms (residues 601 to 627) and the spike core was computed using MDAnalysis^90,91^ to determine flexibility of the 620-loop.

#### Trimeric area calculation for NTD, RBD, RBM, SD2, FPPR

MDAnalysis^90,91^ was used to calculate distances and areas between trimeric domains of interest in the SHC014 spike protein. The calculation was done by selecting the center of mass of each domain of interest per chain (A, B, and C) and determining the distance between that particular domain per chain, and then calculating area via the three-sided area formula. These calculations were performed per frame over all frames in all simulations, and per variant. The residue numbers for each domain are as follows: NTD (18 to 279), RBD (319 to 515), RBM (424 to 495), SD2 (577 to 680), and FPPR (818 to 838).

#### Accessible surface area (ASA)

SASA was computed using the measure sasa command in VMD^58^, which implements the Shrake and Rupley method, with in-house Tcl scripts. A probe radii 7.2Å was used to assess surface accessibility of the SHC014 spike protein. To estimate glycan shielding, the ASA of the protein with glycans was subtracted from that of the same protein lacking glycans. These calculations were carried out at 1-nanosecond intervals across the trajectories, and the results were averaged over all replicas, with their respective standard deviations.

### Dynamical Network Analysis using WISP

To understand the allosteric communication that may influence conformational changes in RBD of SHC014 spike variants, we employed a dynamical network analysis approach using data from our conventional MD simulations. Then, we computed distance and cross-correlation matrices using the cpptraj module of Amber^92,93^, considering the Cα atoms of protein residues. The cross-correlation matrix, calculated using Pearson’s correlation coefficient, quantifies residue pair correlations by measuring the linear relationship between atomic motions. The correlation coefficient values range from -1 to 1, where +1 indicates residues moving in the same direction (fully correlated motion), 0 represents uncorrelated motion, and -1 denotes residues moving in opposite directions (fully anti-correlated motion).

By utilizing the Weighted Implementation of Suboptimal Paths (WISP) method^68^, we constructed a network representation of the Spike protein by mapping the N residues into N nodes of a weighted graph. Each residue was represented as a node, centered on its Cα, and an edge was established between two residues with a mean distance cutoff of 6 Å throughout the simulation. The edges were weighted based on the normalized cross-correlation matrix values using the transformation (dij=-log |Cij|), where Cij is the cross-correlation coefficient between residues i and j, and dij represents a functional correlation-based distance. This transformation ensures that strongly correlated residue pairs have shorter edge distances, reflecting the probability of information transfer based on their correlation. Dijkstra’s algorithm^94^, implemented in the NetworkX Python Library, was employed to uncover critical allosteric communication pathways. This algorithm determines the shortest paths between residues by ranking edges with shorter distances, indicating the most frequently used and highly correlated pathways. The method identifies the highest-contributing edges to allosteric communication and the residues that are key to network connectivity. The top 1000 edges and nodes from the dynamical networks contributing to the allosteric pathways were visualized using VMD^58^. Finally, we extracted the edge-weighted domain-domain pairs to map continuous pathways and reveal how the F294L and A835D mutations reshape the communication routes.

## Abbreviations

MD: Molecular dynamics
NTD: N-terminal Domain
RBD: Receptor Binding Domain
RBM: Receptor Binding Motif
SD1: Sub Domain 1
SD2: Sub Domain 2
FCS: Furin Cleavage Site
FP: Fusion Peptide
FPPR: Fusion Peptide Proximal Region
HR1: Heptad Repeat 1
CD: Connecting Domain
DM: Double mutant

## Data availability

All simulation trajectories, simulation scripts, analysis scripts, and data files will be made accessible for download on the Amaro Lab website at this link (https://amarolab.ucsd.edu/data.php#covid19)

## Author Contributions

TAB, FLK, CCT, LC, designed and conducted all-atom conventional molecular dynamics simulations. TAB, FLK, CCT performed all computational analyses. ALT, CMA, GL performed all viral assays and CryoEM experimentation in previous study^51^, which served as the basis of this work by identifying the F294L and A835D mutations, and also resolved cryoEM SHC014 WT spike structure (PDB ID: 9CAS) used herein. EHM, KC, and JSM guided all previous experiments^51^, and REA oversaw all simulations and analyses.

## Funding Sources

T.A.B is supported by the Shurl and Kay Curci Foundation (https://curcifoundation.org/), the A.G. Leventis Foundation (https://www.leventisfoundation.org/), and Thermo Fisher Scientific Antibody Fellowships (https://www.thermofisher.com/us/en/home/lifescience/antibodies/thermo-fisherscientificantibody-scholarship-program.html).

## Acknowledgments

We are grateful for the Leadership Resource Allocation (LRAC- CHE23002) to use the Frontera Supercomputer provided by the Texas Advanced Computing Centre (TACC). Additional computing resources were provided by the Triton Shared Computing Cluster (TSCC) at San Diego Supercomputer Centre.

## Declaration of Interests

K.C. is a member of the scientific advisory board and holds shares in Integrum Scientific, LLC. K.C. is a cofounder of and holds shares in Eitr Biologics Inc. All other authors declare no competing interests.

