## Supplementary Information for "Structural Dynamics and Allosteric Communication of a SARS-Like Bat Coronavirus Spike Glycoprotein"

### Contents

**Figure S1:** Modeling of missing regions of SHC014 GP
**Figure S2:** Root mean square deviation (RMSD) of SHC014 variant simulations
**Figure S3:** Trimeric area calculation for NTD, RBD, RBM, SD2 and FPPR
**Figure S4:** RBD trimeric area
**Figure S5:** RBD-core distances
**Figure S6:** Accessible surface area of SHC014 spike domains
**Figure S7:** X294-K266 contacts
**Figure S8:** X294-K288 contacts
**Figure S9:** X294-D275 contacts
**Figure S10:** X835-R830 contacts
**Figure S11:** R830-D555 contacts
**Figure S12:** X835-D555 contacts
**Figure S13:** Salt-bridge formation and A835-R830-D555 area

**Figure S14:** R830-N270 glycan contacts
**Figure S15:** R830-N590 glycan
**Figure S16:** Conformational flexibility of the 620-loop in SHC014 WT and mutants
**Figure S17:** Domain-domain communication

**Table S1:** Complete details of SHC014 computational models and simulation information
**Table S2:** Summary of the full-length S protein all-atom MD simulations
**Table S3.** Glycan compositions for chains A, B, and C.

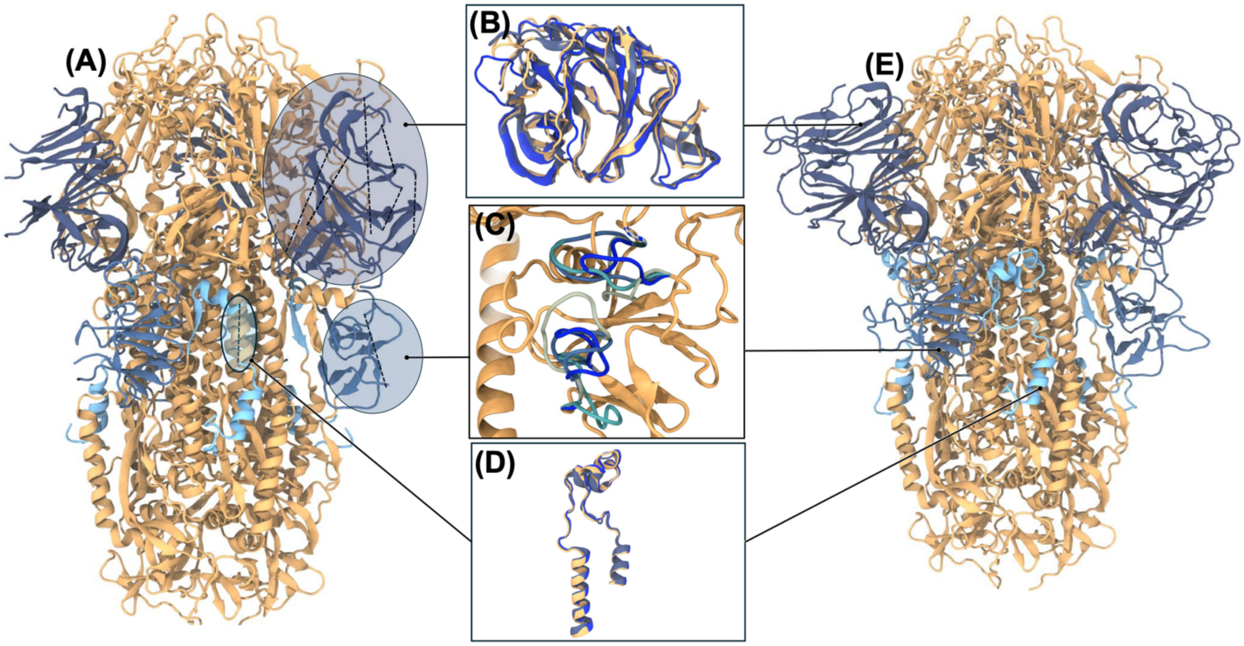


**Figure S1:** Modeling of missing regions of SHC014 GP (A) Wildtype cryoEM structure of SCH01with missing regions highlighted in broken lines with light filled sphere (B) NTD modeling using multiple protein structure prediction tools such as Alphafold (darkblue), Prime (blue) and SwissModel (Gold) (C) Modeling of missing loops in SD2 domain using ChimeraX (D) FPPR modeling using PDB 7JJI as template shown in blue, SwissModel (Gold) and Alphafold (darkblue) (E) SHC014 GP ectodomain with missing regions filled.

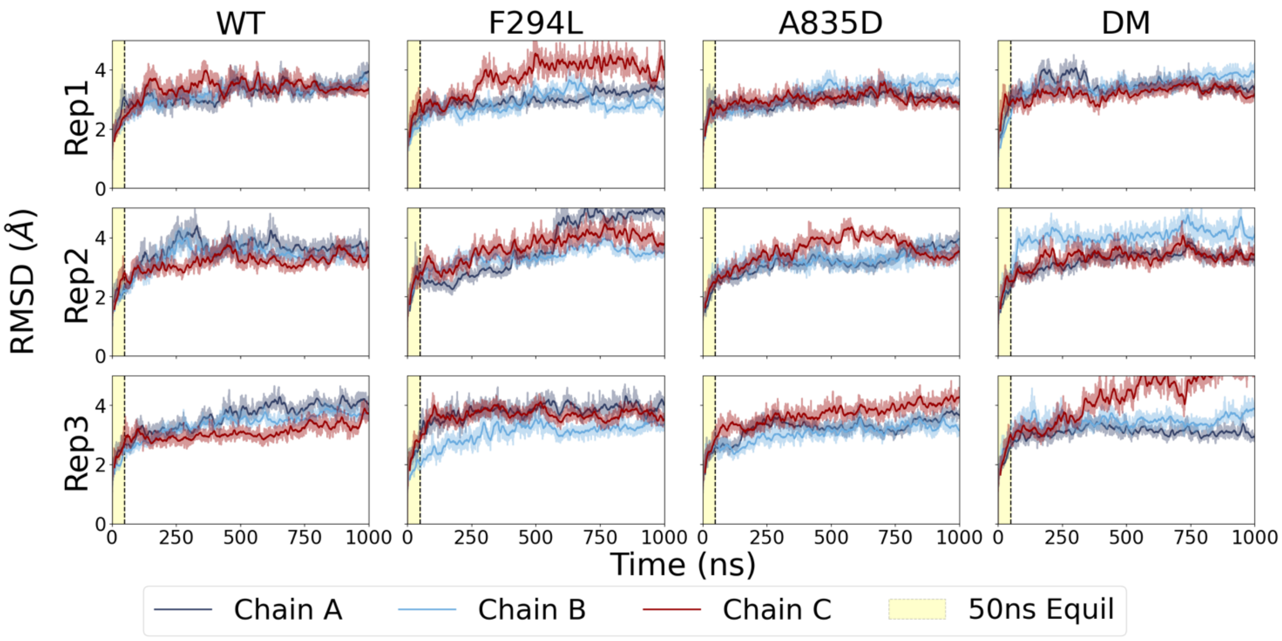


**Figure S2**: Root mean square deviation (RMSD) of SHC014 variants, the 50ns equilibration step before production run is shown in yellow.

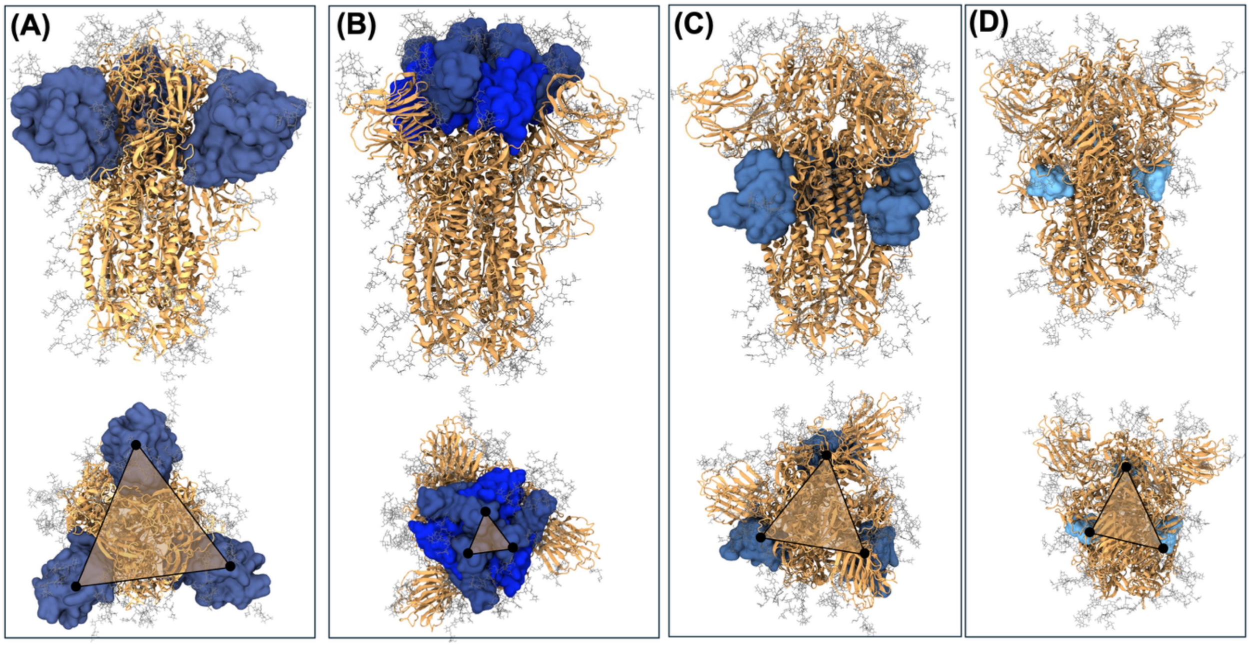


**Figure S3:** Trimeric area calculations of (A) NTD; N-terminal domain (B) RBM; Receptor binding domain (C) SD2; Sub-domain 2 (D) FPPR; Fusion Peptide Proximal Region

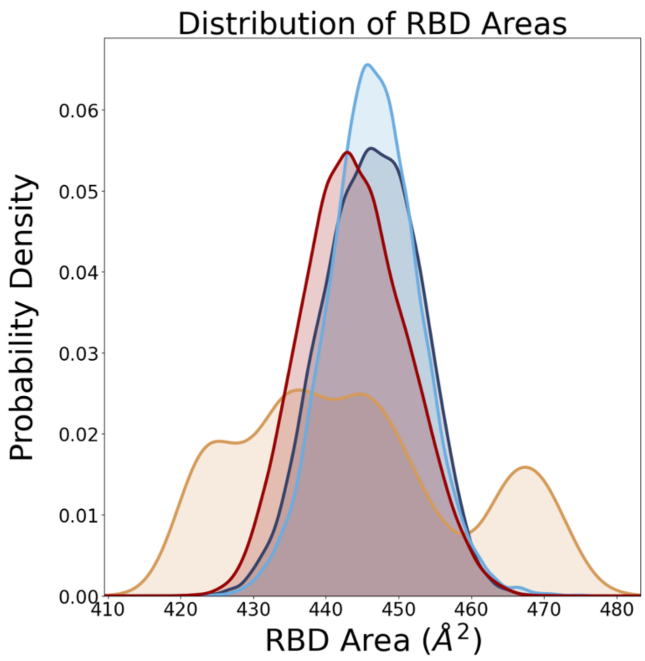


**Figure S4:** RBD trimeric area over the course of the simulations. WT is shown in gold, F294L in dark blue, A835D in cyan, and DM in red.

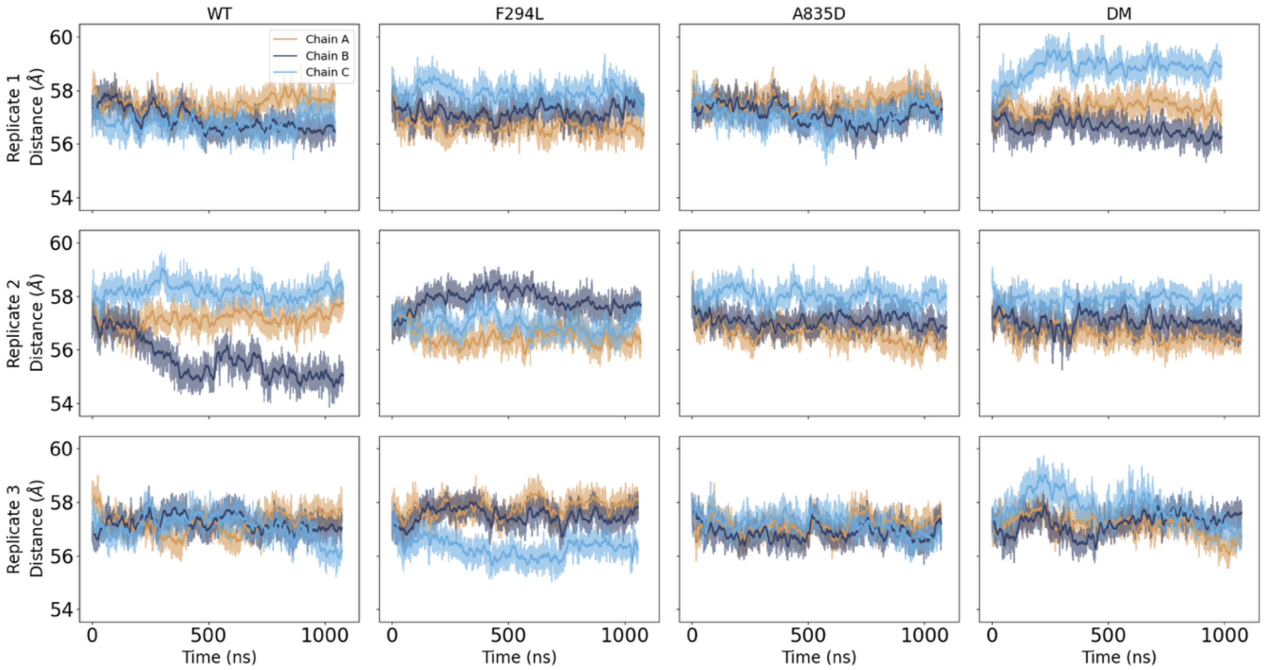


**Figure S5:** Time series analysis of RBD-core distances per replica across the variants depicting the propensity of RBD opening in each chain. Chain A is shown in gold, chain B in dark blue and chain C in cyan.

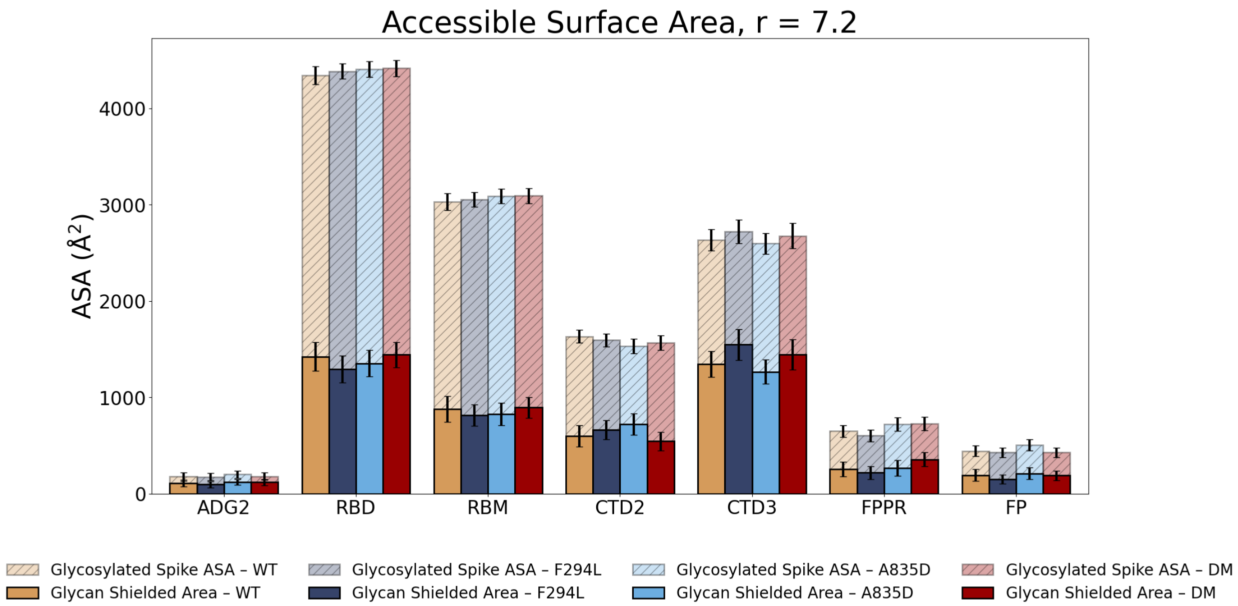


**Figure S6.** Accessible surface area of SHC014 spike domains with a probe radius of 7.2Å. ADG2 referred to the ADG2 binding site.

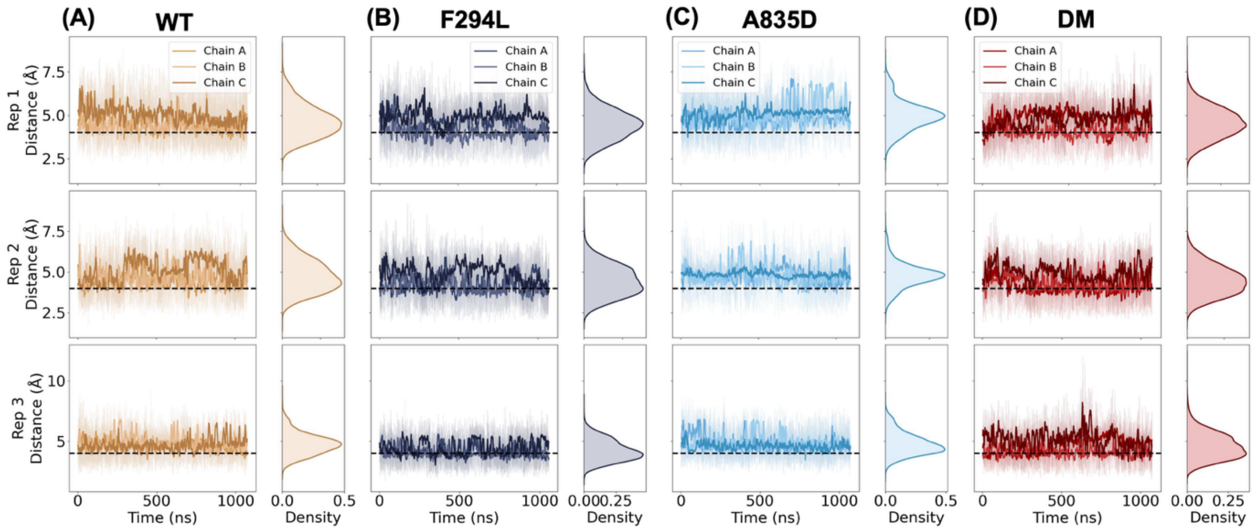


**Figure S7:** X294-K266 contact analysis across the variants with a distance threshold of 4Å shown as dashed lines.

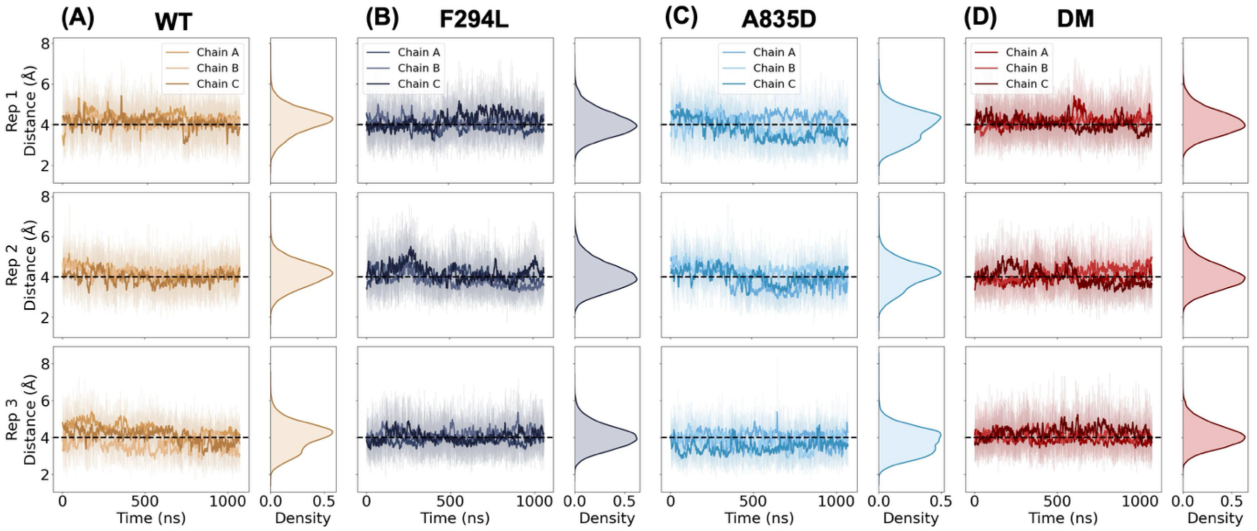


**Figure S8:** X294-K288 contact analysis across the variants with a distance threshold of 4Å shown as dashed lines

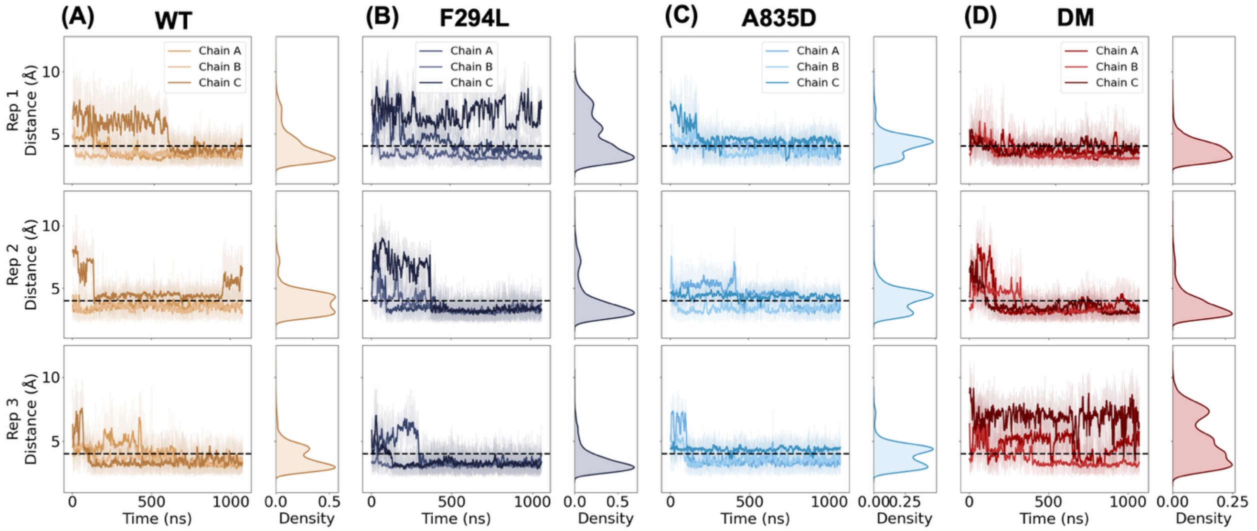


**Figure S9:** X294-D275 contact analysis across the variants with a distance threshold of 4Å shown as dashed lines.

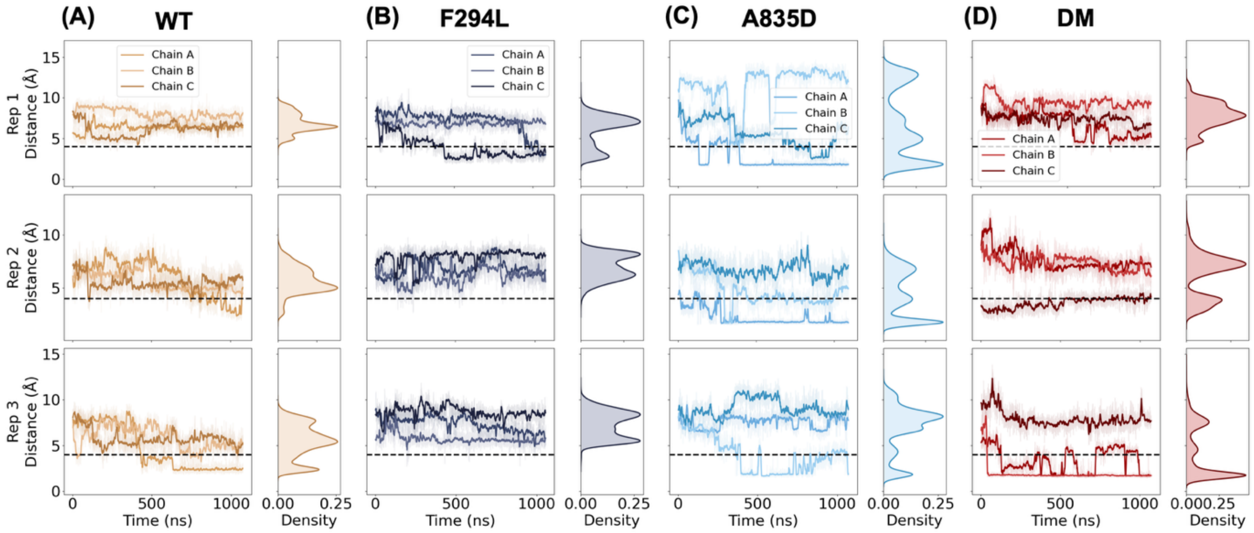


**Figure S10:** X835-R830 contact analysis across the variants with a distance threshold of 4Å shown as dashed lines.

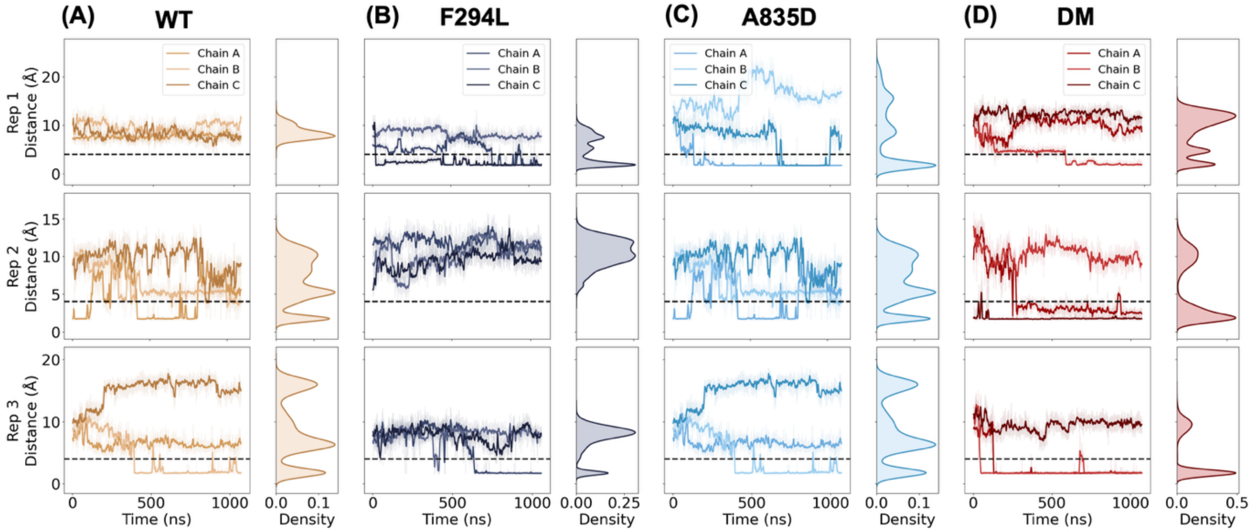


**Figure S11:** R830-D555 contact analysis across the variants with a distance threshold of 4Å shown as dashed lines.

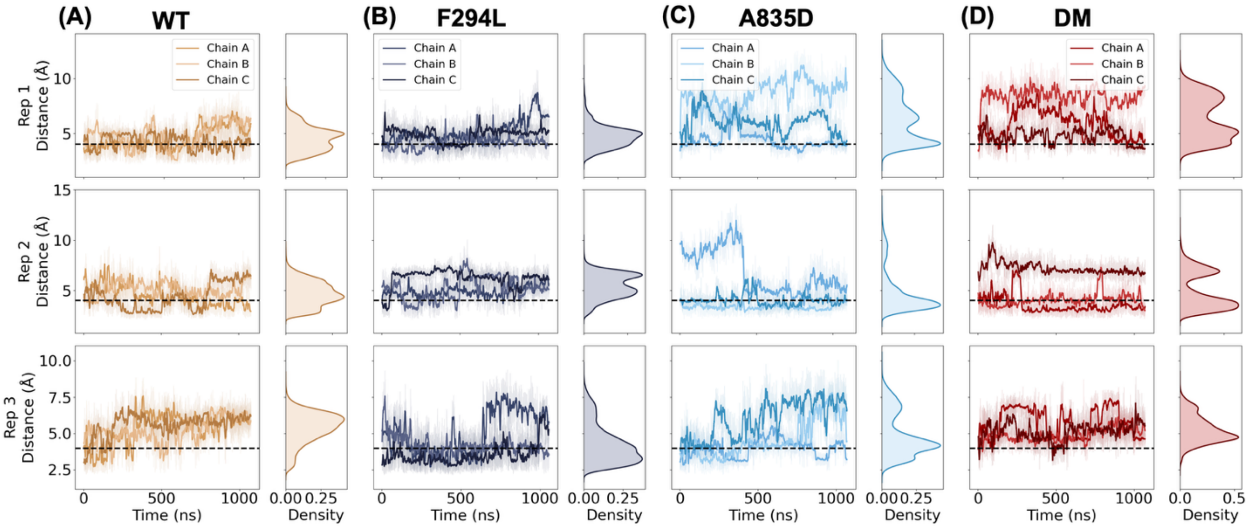


**Figure S12:** X835-D555 contact analysis across the variants with a distance threshold of 4Å shown as dashed lines.

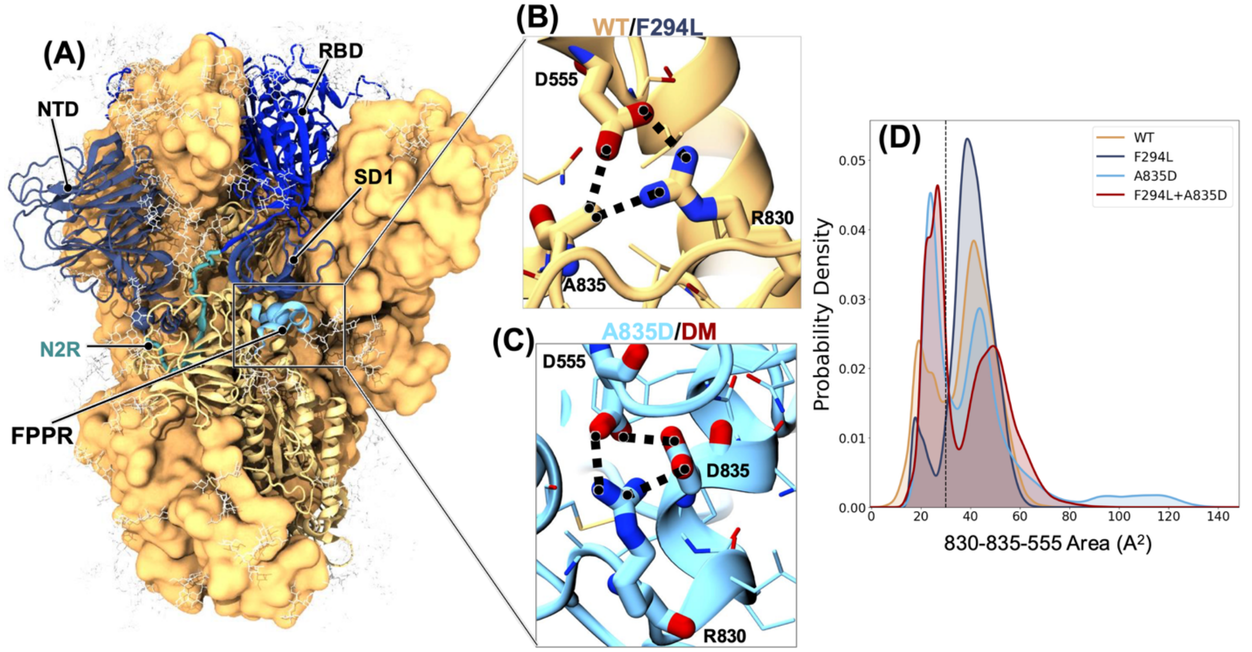


**Figure S13.** Salt bridge formation (A) Highlight of different domains in the SHC014 spike ectodomain, which includes the NTD: N-terminal domain; RBD: Receptor binding domain; SD1: Subdomain-1; N2R Linker and FPPR: Fusion peptide proximal region. Position 835 is shown as a cyan sphere matching the A835D color (B). Close view of A835-R830-D555 area (C) Close view of D835-R830-D555 area (D). Formation and persistence of the salt-bridge triad between R830-D555-X835 with a tight salt-bridge having a threshold of <30Å$^2$ shown as a broken line.

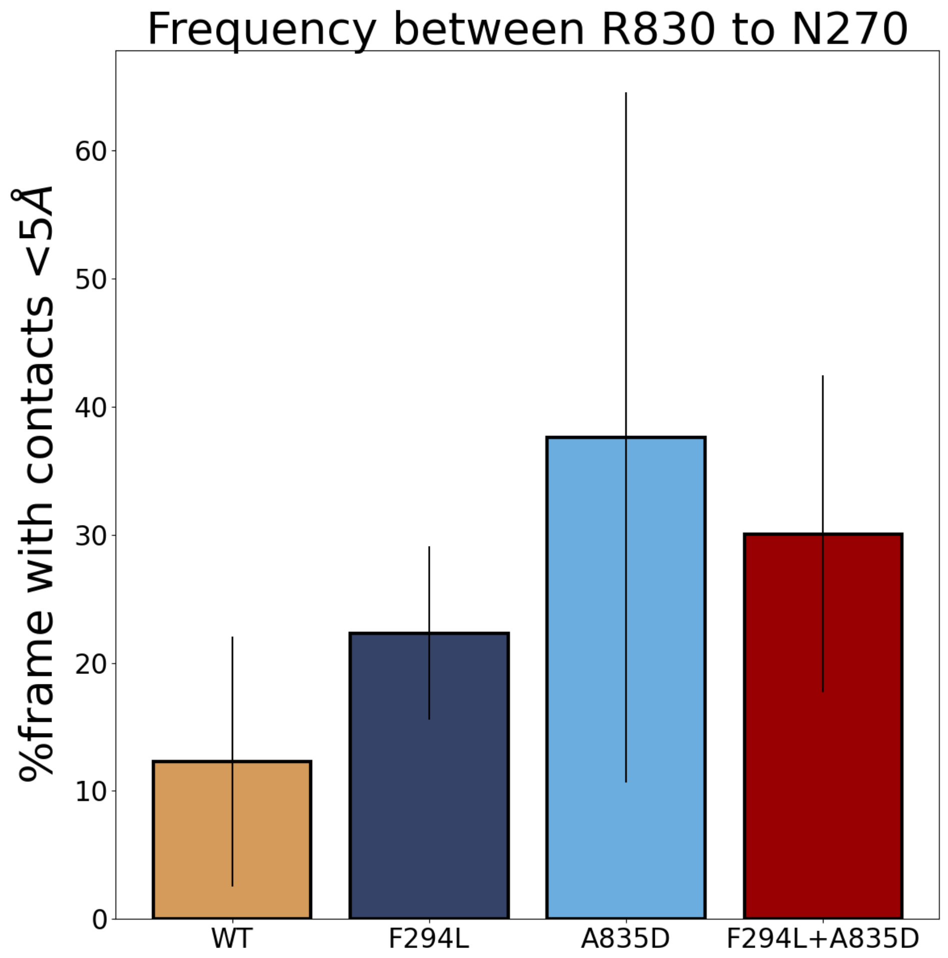


**Figure S14.** Frequency of contact across variants between R830 and glycan N270.

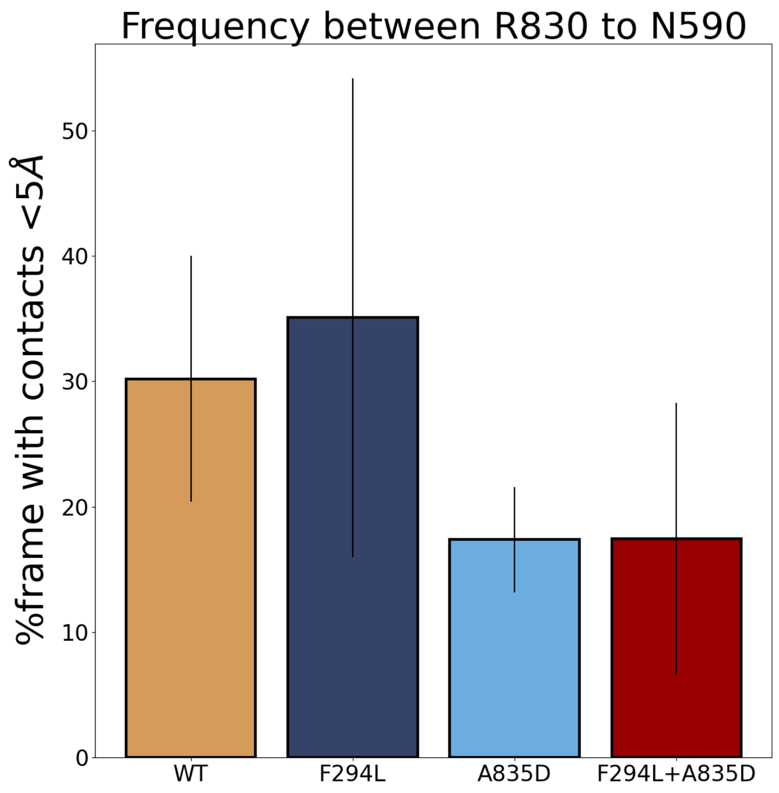


**Figure S15.** Frequency of contact across variants between R830 and glycan N590.

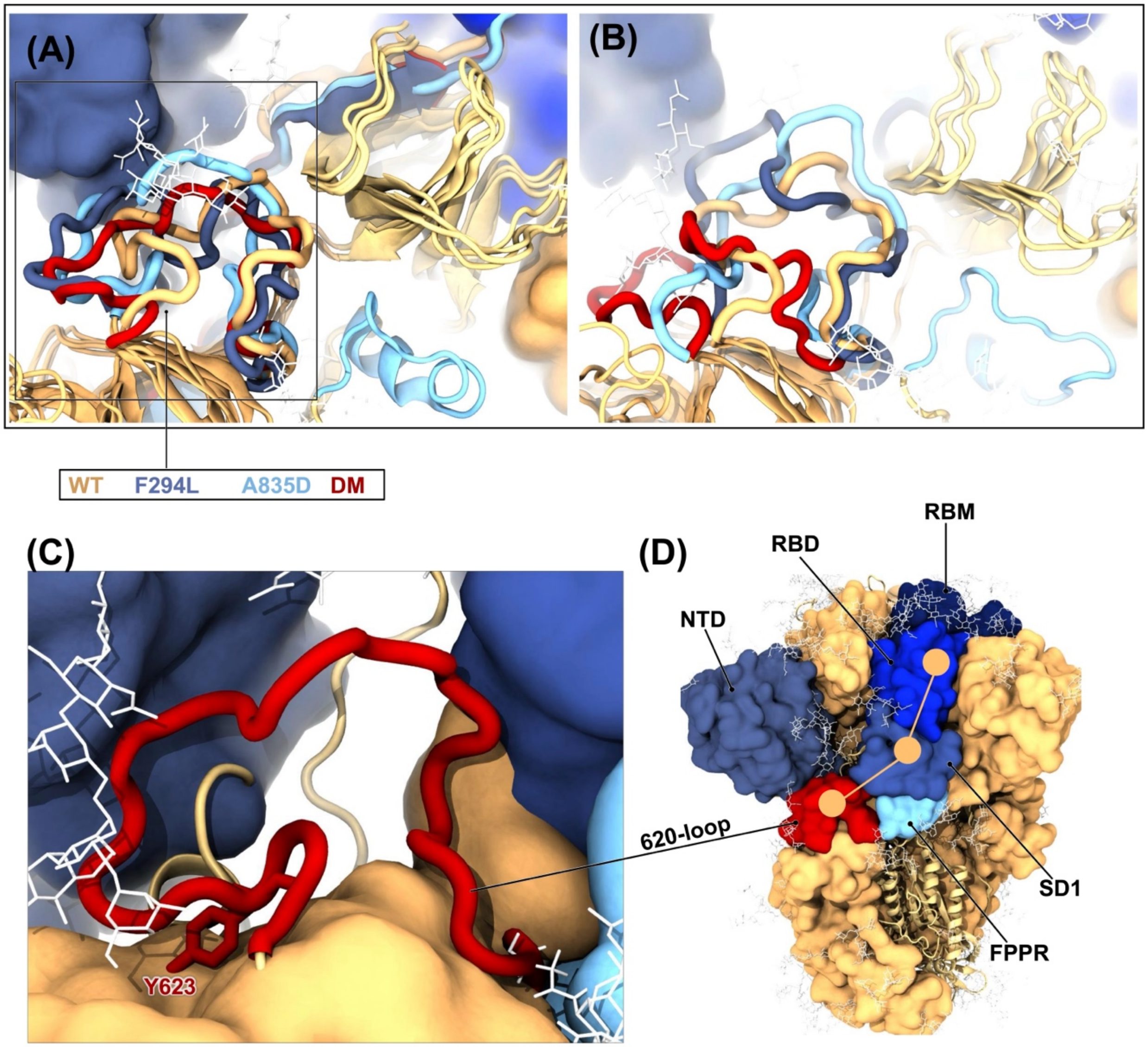


**Figure S16.** Conformational flexibility of the 620-loop across the variants (A). Similar conformation of the 620-loop across the variants (B). Varying degrees in the conformational dynamics of the 620-loop with the double mutant (shown in red), 620-loop showing the highest degree of flexibility (C). A close view of Y623 within the 620loop (D). A proposed model of allosteric communication is depicted by connecting circles from the 620loop through sub-domain-1 to the RBD.

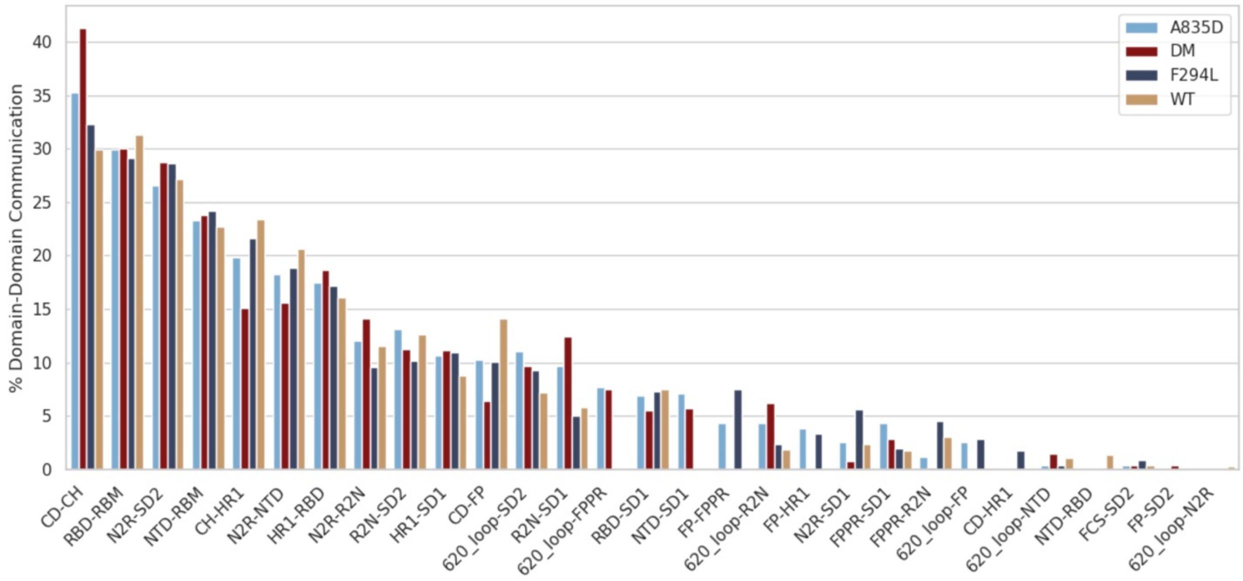


**Figure S17.** Domain-domain communication across the SHC014 spike variants.

**Supplementary Tables**

**Table S1:** Complete details of SHC014 computational models and simulation information:

| | **WT** | **F294L** | **A835D** | **DM** |
|---|---|---|---|---|
| Total #atoms | 972591 | 972585 | 972600 | 972588 |
| #protein atoms/residues | 50928 / 308316 | 50925 / 308315 | 50934 / 308319 | 50931 / 308316 |
| #glycan atoms/glycans | 11415 / 57 | 11415 / 57 | 11415 / 57 | 11415 / 57 |
| #water atoms/residues | 908598 / 302866 | 908595 / 302865 | 908598 / 302866 | 908589 / 302863 |
| #Na/Cl ions | 843 / 807 | 843 / 807 | 846 / 807 | 846 / 807 |
| Disulfides per monomer/ Total disulfide bonds | 15 / 45 | 15 / 45 | 15 / 45 | 15/45 |
| System Dimensions (ÅxÅxÅ) | 215.00 X 214.99 X 214.98 | 215.00 X 214.99 X 214.98 | 215.00 X 214.99 X 214.98 | 215.00 X 214.99 X 214.98 |

**Table S2:** Summary of the full-length S protein all-atom MD simulations.

| **Variant** | **Rep1 (ns)** | **Rep2 (ns)** | **Rep3 (ns)** |
|---|---|---|---|
| WT | 1042.3 | 1078.3 | 1074.2 |
| F294L | 1080.3 | 1068.8 | 1056.8 |
| A835D | 1073.8 | 1093.2 | 1072 |
| DM | 1075.6 | 1072 | 1073.1 |

**Table S3.** Glycan compositions for chains A, B, and C.

**CHAIN A**

| # | SITE | TYPE | STRUCTURE | SEQUENCE |
|---|---|---|---|---|
| G1 | N66 | M5 | 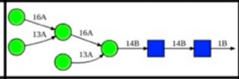 | aDMan(1→6)[aDMan(1→3)]aDMan(1→6)<br>[aDMan(1→3)]bDMan(1→4)bDGlcNAc(1→4)bDGlcNAc(1→)PROA-66 |
| G2 | N110 | FA2G2S1 | 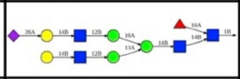 | aDNeu5Ac(2→6)bDGal(1→4)bDGlcNAc(1→2)aDMan(1→6)<br>[bDGal(1→4)bDGlcNAc(1→2)aDMan(1→3)]bDMan(1→4)bDGlcNAc(1→4)<br>[aLFuc(1→6)]bDGlcNAc(1→)PROA-110 |
| G3 | N120 | M5 | 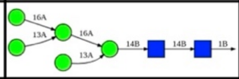 | aDMan(1→6)[aDMan(1→3)]aDMan(1→6)<br>[aDMan(1→3)]bDMan(1→4)bDGlcNAc(1→4)bDGlcNAc(1→)PROA-120 |
| G4 | N145 | FA2G2S1 | 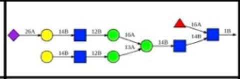 | aDNeu5Ac(2→6)bDGal(1→4)bDGlcNAc(1→2)aDMan(1→6)<br>[bDGal(1→4)bDGlcNAc(1→2)aDMan(1→3)]bDMan(1→4)bDGlcNAc(1→4)<br>[aLFuc(1→6)]bDGlcNAc(1→)PROA-145 |
| G5 | N159 | FA2G2S2 | 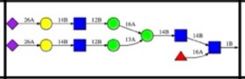 | aDNeu5Ac(2→6)bDGal(1→4)bDGlcNAc(1→2)aDMan(1→6)<br>[aDNeu5Ac(2→6)bDGal(1→4)bDGlcNAc(1→2)aDMan(1→3)]bDMan(1→4)bDGlcNAc(1→4)<br>[aLFuc(1→6)]bDGlcNAc(1→)PROA-159 |
| G6 | N228 | M8 | 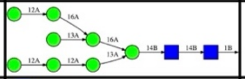 | aDMan(1→2)aDMan(1→6)[aDMan(1→3)]aDMan(1→6)<br>[aDMan(1→2)aDMan(1→2)aDMan(1→3)]bDMan(1→4)bDGlcNAc(1→4)bDGlcNAc(1→)PROA-228 |
| G7 | N270 | FA3 | 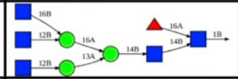 | bDGlcNAc(1→6)[bDGlcNAc(1→2)]aDMan(1→6)<br>[bDGlcNAc(1→2)aDMan(1→3)]bDMan(1→4)bDGlcNAc(1→4)[aLFuc(1→6)]bDGlcNAc(1→)PROA-270 |
| G8 | N319 | FA2 | 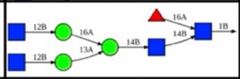 | bDGlcNAc(1→2)aDMan(1→6)[bDGlcNAc(1→2)aDMan(1→3)]bDMan(1→4)bDGlcNAc(1→4)<br>[aLFuc(1→6)]bDGlcNAc(1→)PROA-319 |
| G9 | N331 | FA2 | 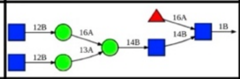 | bDGlcNAc(1→2)aDMan(1→6)[bDGlcNAc(1→2)aDMan(1→3)]bDMan(1→4)bDGlcNAc(1→4)<br>[aLFuc(1→6)]bDGlcNAc(1→)PROA-331 |
| G10 | N358 | FA2G2S1 | 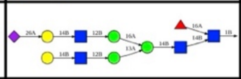 | aDNeu5Ac(2→6)bDGal(1→4)bDGlcNAc(1→2)aDMan(1→6)<br>[bDGal(1→4)bDGlcNAc(1→2)aDMan(1→3)]bDMan(1→4)bDGlcNAc(1→4)<br>[aLFuc(1→6)]bDGlcNAc(1→)PROA-358 |
| G11 | N590 | FA2 | 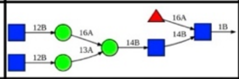 | bDGlcNAc(1→2)aDMan(1→6)[bDGlcNAc(1→2)aDMan(1→3)]bDMan(1→4)bDGlcNAc(1→4)<br>[aLFuc(1→6)]bDGlcNAc(1→)PROA-590 |
| G12 | N603 | A2 | 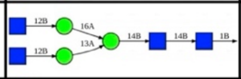 | bDGlcNAc(1→2)aDMan(1→6)<br>[bDGlcNAc(1→2)aDMan(1→3)]bDMan(1→4)bDGlcNAc(1→4)bDGlcNAc(1→)PROA-603 |
| G13 | N692 | M6 | 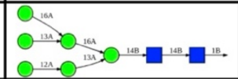 | aDMan(1→6)[aDMan(1→3)]aDMan(1→6)<br>[aDMan(1→2)aDMan(1→3)]bDMan(1→4)bDGlcNAc(1→4)bDGlcNAc(1→)PROA-692 |
| G14 | N700 | Hybrid G1 | 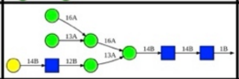 | bDGal(1→4)bDGlcNAc(1→2)aDMan(1→3)[aDMan(1→6)<br>[aDMan(1→3)]aDMan(1→6)]bDMan(1→4)bDGlcNAc(1→4)bDGlcNAc(1→)PROA-700 |
| G15 | N784 | M6 | 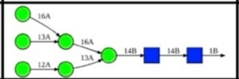 | aDMan(1→6)[aDMan(1→3)]aDMan(1→6)<br>[aDMan(1→2)aDMan(1→3)]bDMan(1→4)bDGlcNAc(1→4)bDGlcNAc(1→)PROA-784 |
| G16 | N1057 | FA2G2S1 | 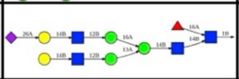 | aDNeu5Ac(2→6)bDGal(1→4)bDGlcNAc(1→2)aDMan(1→6)<br>[bDGal(1→4)bDGlcNAc(1→2)aDMan(1→3)]bDMan(1→4)bDGlcNAc(1→4)<br>[aLFuc(1→6)]bDGlcNAc(1→)PROA-1057 |
| G17 | N1081 | FA2 | 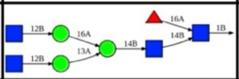 | bDGlcNAc(1→2)aDMan(1→6)[bDGlcNAc(1→2)aDMan(1→3)]bDMan(1→4)bDGlcNAc(1→4)<br>[aLFuc(1→6)]bDGlcNAc(1→)PROA-1081 |
| G18 | N1117 | FA1 | 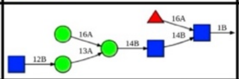 | bDGlcNAc(1→2)aDMan(1→3)[aDMan(1→6)]bDMan(1→4)bDGlcNAc(1→4)<br>[aLFuc(1→6)]bDGlcNAc(1→)PROB-1117 |
| G19 | S311 | O-glycan | 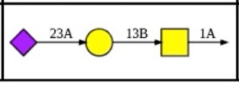 | aDNeu5Ac(2→3)bDGal(1→3)aDGalNAc(1→)PROA-311 |

CHAIN B

| # | SITE | TYPE | STRUCTURE | SEQUENCE |
|---|---|---|---|---|
| G20 | N66 | M5 | 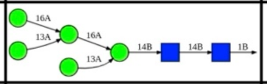 | aDMan(1→6)[aDMan(1→3)]aDMan(1→6)<br>[aDMan(1→3)]bDMan(1→4)bDGlcNAc(1→4)bDGlcNAc(1→)PROB-66 |
| G21 | N110 | FA2G2S2 | 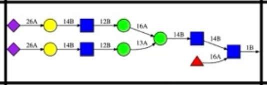 | aDNeu5Ac(2→6)bDGal(1→4)bDGlcNAc(1→2)aDMan(1→6)<br>[aDNeu5Ac(2→6)bDGal(1→4)bDGlcNAc(1→2)aDMan(1→3)]bDMan(1→4)bDGlcNAc(1→4)<br>[aLFuc(1→6)]bDGlcNAc(1→)PROB-110 |
| G22 | N120 | FA2 | 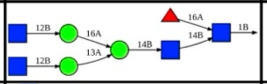 | bDGlcNAc(1→2)aDMan(1→6)[bDGlcNAc(1→2)aDMan(1→3)]bDMan(1→4)bDGlcNAc(1→4)<br>[aLFuc(1→6)]bDGlcNAc(1→)PROB-120 |
| G23 | N145 | FA3 | 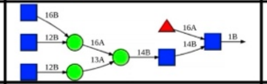 | bDGlcNAc(1→6)[bDGlcNAc(1→2)]aDMan(1→6)<br>[bDGlcNAc(1→2)aDMan(1→3)]bDMan(1→4)bDGlcNAc(1→4)[aLFuc(1→6)]bDGlcNAc(1→)PROB-145 |
| G24 | N159 | M5 | 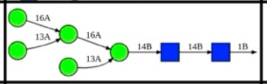 | aDMan(1→6)[aDMan(1→3)]aDMan(1→6)<br>[aDMan(1→3)]bDMan(1→4)bDGlcNAc(1→4)bDGlcNAc(1→)PROB-159 |
| G25 | N228 | M9 | 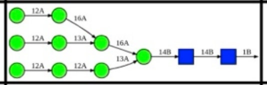 | aDMan(1→2)aDMan(1→6)[aDMan(1→2)aDMan(1→3)]aDMan(1→6)<br>[aDMan(1→2)aDMan(1→2)aDMan(1→3)]bDMan(1→4)bDGlcNAc(1→4)bDGlcNAc(1→)PROB-228 |
| G26 | N270 | FA3G3S1 | 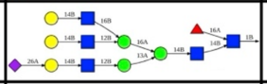 | aDNeu5Ac(2→6)bDGal(1→4)bDGlcNAc(1→2)aDMan(1→3)[bDGal(1→4)bDGlcNAc(1→6)<br>[bDGal(1→4)bDGlcNAc(1→2)]aDMan(1→6)]bDMan(1→4)bDGlcNAc(1→4)<br>[aLFuc(1→6)]bDGlcNAc(1→)PROB-270 |
| G27 | N319 | FA2 | 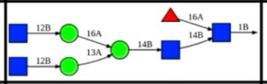 | bDGlcNAc(1→2)aDMan(1→6)[bDGlcNAc(1→2)aDMan(1→3)]bDMan(1→4)bDGlcNAc(1→4)<br>[aLFuc(1→6)]bDGlcNAc(1→)PROB-319 |
| G28 | N331 | FA1 | 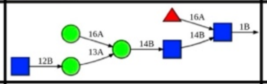 | bDGlcNAc(1→2)aDMan(1→3)[aDMan(1→6)]bDMan(1→4)bDGlcNAc(1→4)<br>[aLFuc(1→6)]bDGlcNAc(1→)PROB-331 |
| G29 | N358 | FA2G2S2 | 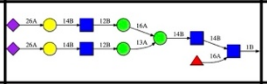 | aDNeu5Ac(2→6)bDGal(1→4)bDGlcNAc(1→2)aDMan(1→6)<br>[aDNeu5Ac(2→6)bDGal(1→4)bDGlcNAc(1→2)aDMan(1→3)]bDMan(1→4)bDGlcNAc(1→4)<br>[aLFuc(1→6)]bDGlcNAc(1→)PROB-358 |
| G30 | N590 | M5 | 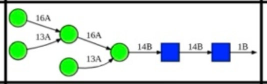 | aDMan(1→6)[aDMan(1→3)]aDMan(1→6)<br>[aDMan(1→3)]bDMan(1→4)bDGlcNAc(1→4)bDGlcNAc(1→)PROB-590 |
| G31 | N603 | FA2 | 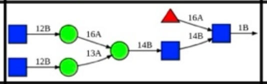 | bDGlcNAc(1→2)aDMan(1→6)[bDGlcNAc(1→2)aDMan(1→3)]bDMan(1→4)bDGlcNAc(1→4)<br>[aLFuc(1→6)]bDGlcNAc(1→)PROB-603 |
| G32 | N692 | M5 | 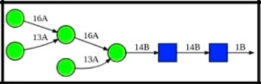 | aDMan(1→6)[aDMan(1→3)]aDMan(1→6)<br>[aDMan(1→3)]bDMan(1→4)bDGlcNAc(1→4)bDGlcNAc(1→)PROB-692 |
| G33 | N700 | M5 | 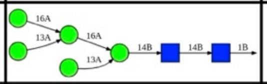 | aDMan(1→6)[aDMan(1→3)]aDMan(1→6)<br>[aDMan(1→3)]bDMan(1→4)bDGlcNAc(1→4)bDGlcNAc(1→)PROB-700 |
| G34 | N784 | M7 | 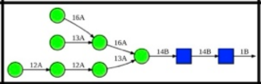 | aDMan(1→2)aDMan(1→2)aDMan(1→3)[aDMan(1→6)<br>[aDMan(1→3)]aDMan(1→6)]bDMan(1→4)bDGlcNAc(1→4)bDGlcNAc(1→)PROB-784 |
| G35 | N1057 | M5 | 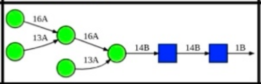 | aDMan(1→6)[aDMan(1→3)]aDMan(1→6)<br>[aDMan(1→3)]bDMan(1→4)bDGlcNAc(1→4)bDGlcNAc(1→)PROB-1057 |
| G36 | N1081 | A2 | 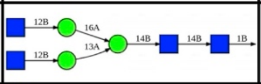 | bDGlcNAc(1→2)aDMan(1→6)<br>[bDGlcNAc(1→2)aDMan(1→3)]bDMan(1→4)bDGlcNAc(1→4)bDGlcNAc(1→)PROB-1081 |
| G37 | N1117 | FA3 | 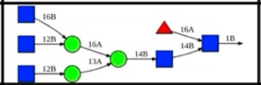 | bDGlcNAc(1→6)[bDGlcNAc(1→2)]aDMan(1→6)<br>[bDGlcNAc(1→2)aDMan(1→3)]bDMan(1→4)bDGlcNAc(1→4)[aLFuc(1→6)]bDGlcNAc(1→)PROB-1117 |
| G38 | S311 | O-glycan | 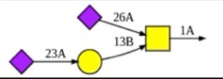 | aDNeu5Ac(2→3)bDGal(1→3)[aDNeu5Ac(2→6)]aDGalNAc(1→)PROB-311 |

CHAIN C

| # | SITE | TYPE | STRUCTURE | SEQUENCE |
|---|---|---|---|---|
| G39 | N66 | M5 | 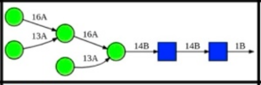 | aDMan(1→6)[aDMan(1→3)]aDMan(1→6)<br>[aDMan(1→3)]bDMan(1→4)bDGlcNAc(1→4)bDGlcNAc(1→)PROC-66 |
| G40 | N110 | M8 | 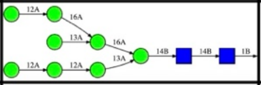 | aDMan(1→2)aDMan(1→6)[aDMan(1→3)]aDMan(1→6)<br>[aDMan(1→2)aDMan(1→2)aDMan(1→3)]bDMan(1→4)bDGlcNAc(1→4)bDGlcNAc(1→)PROC-110 |
| G41 | N120 | M5 | 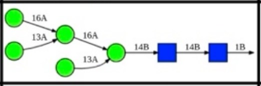 | aDMan(1→6)[aDMan(1→3)]aDMan(1→6)<br>[aDMan(1→3)]bDMan(1→4)bDGlcNAc(1→4)bDGlcNAc(1→)PROC-120 |
| G42 | N145 | FA2 | 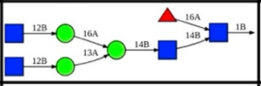 | bDGlcNAc(1→2)aDMan(1→6)[bDGlcNAc(1→2)aDMan(1→3)]bDMan(1→4)bDGlcNAc(1→4)<br>[aLFuc(1→6)]bDGlcNAc(1→)PROC-145 |
| G43 | N159 | FA2G2S1 | 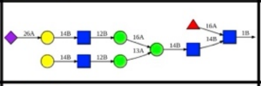 | aDNeu5Ac(2→6)bDGal(1→4)bDGlcNAc(1→2)aDMan(1→6)<br>[bDGal(1→4)bDGlcNAc(1→2)aDMan(1→3)]bDMan(1→4)bDGlcNAc(1→4)<br>[aLFuc(1→6)]bDGlcNAc(1→)PROC-159 |
| G44 | N228 | M9 | 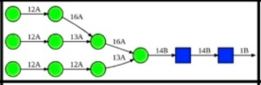 | aDMan(1→2)aDMan(1→6)[aDMan(1→2)aDMan(1→3)]aDMan(1→6)<br>[aDMan(1→2)aDMan(1→2)aDMan(1→3)]bDMan(1→4)bDGlcNAc(1→4)bDGlcNAc(1→)PROC-228 |
| G45 | N270 | A2 | 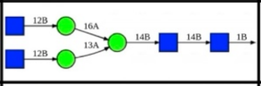 | bDGlcNAc(1→2)aDMan(1→6)<br>[bDGlcNAc(1→2)aDMan(1→3)]bDMan(1→4)bDGlcNAc(1→4)bDGlcNAc(1→)PROC-270 |
| G46 | N319 | FA3G3S1 | 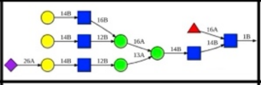 | aDNeu5Ac(2→6)bDGal(1→4)bDGlcNAc(1→2)aDMan(1→3)[bDGal(1→4)bDGlcNAc(1→6)<br>[bDGal(1→4)bDGlcNAc(1→2)]aDMan(1→6)]bDMan(1→4)bDGlcNAc(1→4)<br>[aLFuc(1→6)]bDGlcNAc(1→)PROC-319 |
| G47 | N331 | FA2 | 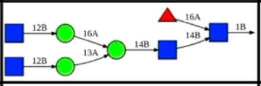 | bDGlcNAc(1→2)aDMan(1→6)[bDGlcNAc(1→2)aDMan(1→3)]bDMan(1→4)bDGlcNAc(1→4)<br>[aLFuc(1→6)]bDGlcNAc(1→)PROC-331 |
| G48 | N358 | FA3G3S1 | 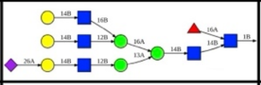 | aDNeu5Ac(2→6)bDGal(1→4)bDGlcNAc(1→2)aDMan(1→3)[bDGal(1→4)bDGlcNAc(1→6)<br>[bDGal(1→4)bDGlcNAc(1→2)]aDMan(1→6)]bDMan(1→4)bDGlcNAc(1→4)<br>[aLFuc(1→6)]bDGlcNAc(1→)PROC-358 |
| G49 | N590 | M5 | 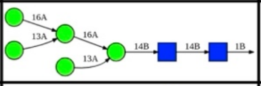 | aDMan(1→6)[aDMan(1→3)]aDMan(1→6)<br>[aDMan(1→3)]bDMan(1→4)bDGlcNAc(1→4)bDGlcNAc(1→)PROC-590 |
| G50 | N603 | FA2 | 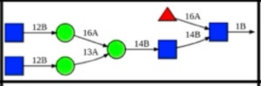 | bDGlcNAc(1→2)aDMan(1→6)[bDGlcNAc(1→2)aDMan(1→3)]bDMan(1→4)bDGlcNAc(1→4)<br>[aLFuc(1→6)]bDGlcNAc(1→)PROC-603 |
| G51 | N692 | M5 | 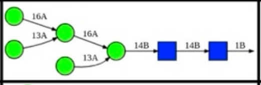 | aDMan(1→6)[aDMan(1→3)]aDMan(1→6)<br>[aDMan(1→3)]bDMan(1→4)bDGlcNAc(1→4)bDGlcNAc(1→)PROC-692 |
| G52 | N700 | M6 | 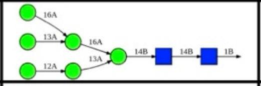 | aDMan(1→6)[aDMan(1→3)]aDMan(1→6)<br>[aDMan(1→2)aDMan(1→3)]bDMan(1→4)bDGlcNAc(1→4)bDGlcNAc(1→)PROC-700 |
| G53 | N784 | M5 | 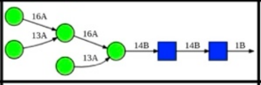 | aDMan(1→6)[aDMan(1→3)]aDMan(1→6)<br>[aDMan(1→3)]bDMan(1→4)bDGlcNAc(1→4)bDGlcNAc(1→)PROC-784 |
| G54 | N1057 | M5 | 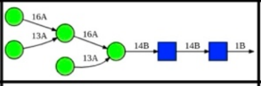 | aDMan(1→6)[aDMan(1→3)]aDMan(1→6)<br>[aDMan(1→3)]bDMan(1→4)bDGlcNAc(1→4)bDGlcNAc(1→)PROC-1057 |
| G55 | N1081 | Hybrid G1S1 | 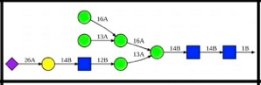 | aDNeu5Ac(2→6)bDGal(1→4)bDGlcNAc(1→2)aDMan(1→3)[aDMan(1→6)<br>[aDMan(1→3)]aDMan(1→6)]bDMan(1→4)bDGlcNAc(1→4)bDGlcNAc(1→)PROC-1081 |
| G56 | N1117 | FA2 | 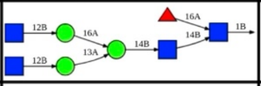 | bDGlcNAc(1→2)aDMan(1→6)[bDGlcNAc(1→2)aDMan(1→3)]bDMan(1→4)bDGlcNAc(1→4)<br>[aLFuc(1→6)]bDGlcNAc(1→)PROC-1117 |
| G57 | S311 | O-glycan | 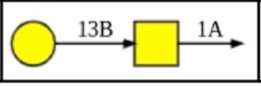 | bDGal(1→3)aDGalNAc(1→)PROC-323 |
